# Neural basis of collective social behavior during environmental challenge

**DOI:** 10.1101/2024.09.17.613378

**Authors:** Tara Raam, Qin Li, Linfan Gu, Gabrielle Elagio, Kayla Y. Lim, Xingjian Zhang, Stephanie M. Correa, Weizhe Hong

## Abstract

Humans and animals have a remarkable capacity to collectively coordinate their behavior to respond to environmental challenges. However, the underlying neurobiology remains poorly understood. Here, we found that groups of mice self-organize into huddles at cold ambient temperature during the thermal challenge assay. We found that mice make active (self-initiated) and passive (partner-initiated) decisions to enter or exit a huddle. Using microendoscopic calcium imaging, we found that active and passive decisions are encoded distinctly within the dorsomedial prefrontal cortex (dmPFC). Silencing dmPFC activity in some mice reduced their active decision-making, but also induced a compensatory increase in active decisions by non-manipulated partners, conserving the group’s overall huddle time. These findings reveal how collective behavior is implemented in neurobiological mechanisms to meet homeostatic needs during environmental challenges.

## MAIN TEXT

Collective behavior, the emergence of group-level dynamics from the actions and interactions of individual animals, is prevalent in many species throughout nature. Collective behaviors such as flocking, schooling, swarming, stampeding, and huddling benefit both the individuals as well as the group as a whole (1,2). In these contexts, each individual receives additional protection from environmental stressors that would not be accessible on their own or even in a dyadic pair (3,4). Environmental challenges such as changes in climate and natural disasters can pose substantial challenges for individuals and groups and the social cohesion afforded by group living can increase the likelihood of survival (5).

Despite the prevalence and importance of collective behavior in nature, we know very little about the underlying neurobiology that enables individuals to coordinate with other members of the group (6). In many collective behaviors, group behavior emerges from individuals adhering to simple, well-defined rules about interaction. However, even though the rules underlying many collective behaviors have been extensively studied, very little is known about the neural mechanisms that enable each individual to implement these rules (7). This is in part because many collective behaviors such as flocking, stampeding, and swarming are difficult to study in laboratory environments and in constrained set-ups necessary to carry out neural recordings and manipulation. Additionally, recent advancements in computer vision tools for automatically tracking the identity and posture of each individual animal during behavior have circumvented technical challenges with studying collective behaviors in laboratory settings (8–10). Although recent studies have made significant headway into identifying the neural mechanisms underlying schooling in fish (11,12), group spatial and acoustic behavior in bats (13,14), and collective escape and defense in flies (15,16), much is still unknown about how groups of animals sense changes in the environment and appropriately coordinate their behavior together.

Here, we use the thermal challenge assay to identify the neural basis of collective huddling behavior at cold ambient temperature in groups of mice. Exposure to cold temperature below thermoneutrality is a stressor for a wide range of animals, including rodents (17). In many endothermic species, exposure to cold temperature can lead to rapid heat loss, triggering increased metabolic rate to sustain core body temperature (18,19). Further, prolonged exposure to cold temperature increases anxiety-like behavior and serum corticosterone (20,21), and can impair gut motility (22) and increase tumorigenesis (23). To mitigate these adverse effects, animals can use a variety of behavioral strategies to thermoregulate such as seeking heat (24) and increasing food consumption (25,26). Notably, animals living in groups such as rodents (27–33), primates (34,35), and penguins (36,37), also organize into collective huddles to thermoregulate.

Huddling behavior shares many features of collective behavior in a variety of species, including emergence, phase transitions below critical temperatures (38), and self-organization without instruction (27,39). However, the underlying neurobiology that enables groups to coordinate into huddles has not been identified. We characterize a novel decision-making framework to identify active (self-initiated) and passive (partner-initiated) decisions to enter and exit huddles. We identify the dorsomedial prefrontal cortex (dmPFC) as an important locus for regulating decisions to engage in huddling. Using microendoscopic calcium imaging, we find that unique populations of neurons in the dmPFC encode active and passive huddling decisons. Moreover, we find that silencing the dmPFC in a group of animals decreases active decisions. Remarkably, when dmPFC is silenced in two out of four animals in the group, we observe that the two non-silenced animals show compensatory changes in their decisions, despite not receiving a direct neural manipulation. Altogether, these findings shed light on the neurobiology of collective behavior and provide new insight into how social groups coordinate their behavior to adapt to environmental challenges.

## RESULTS

### Groups of mice self-organize into huddles in response to cold ambient temperature

To first explore how social groups collectively adapt their behavior in response to environmental demands, we tested groups of male mice in the thermal challenge assay (Fig. 1A). In this task, groups of four co-housed male mice are placed in a temperature-controlled behavior chamber together either at 5°C or 20°C as a control and allowed to freely interact. We found that animals spent a substantial amount of time huddling with each other at 5°C (Fig. 1A-D). To systematically characterize huddling behavior, we developed an automated pipeline to identify the huddling states of the group (Fig. S1). We first used an edge detection function on pixel binarized frames of behavior to identify groups of aggregated animals (Fig. S1A-B). Then, we used SLEAP(8), a deep learning-based neural network used for pose estimation and identity tracking of multiple interacting animals. We combined the outputs of the two algorithms to determine the membership of the huddle (Fig. S1C-D). Using this approach, we identified five unique group huddle states – 0, 2:1:1, 2:2, 3:1, and 4:0, ranging from most dispersed to most aggregated (Fig. 1B). When groups are exposed to cold temperature (5°C), we observed a significant decrease in time that groups spend in the most dispersed state (0 state), and a corresponding increase in time spent in the most aggregated huddle states (3:1 and 4:0) (Fig. 1C). We also observed a decrease in the mean state duration for the 0 state, and an increase in state duration for the aggregated huddle states (Fig. 1D). Interestingly, we found that groups spend very little time in the 2:2 huddle state, suggesting that when all four animals want to huddle, they prefer to do so as a larger group of four where more heat can be generated rather than two smaller groups of two. Notably, the frequency of the states evolved substantially throughout the behavior session. We found that at 5°C, the 0 and 2:1:1 states had a higher frequency early on in the session, but decreased to nearly 0% by the end of the session (Fig. 1E). Conversely, the 3:1:1 and 4:0 states had a low frequency initially, but became the dominant states by the end of the session (Fig. 1E). This suggests that at 5°C, group states evolve dynamically over time, whereas at 20°C, frequency of huddle states is largely stable (Fig. S2F). We also examined the behavior of animal groups under intermediate temperatures of 10°C and 15°C and found that groups huddled more than they did at 20°C, but less than they did at 5°C (Fig. S2), suggesting that huddling behavior is also shaped by the intensity of the thermal challenge.

**Figure 1:**
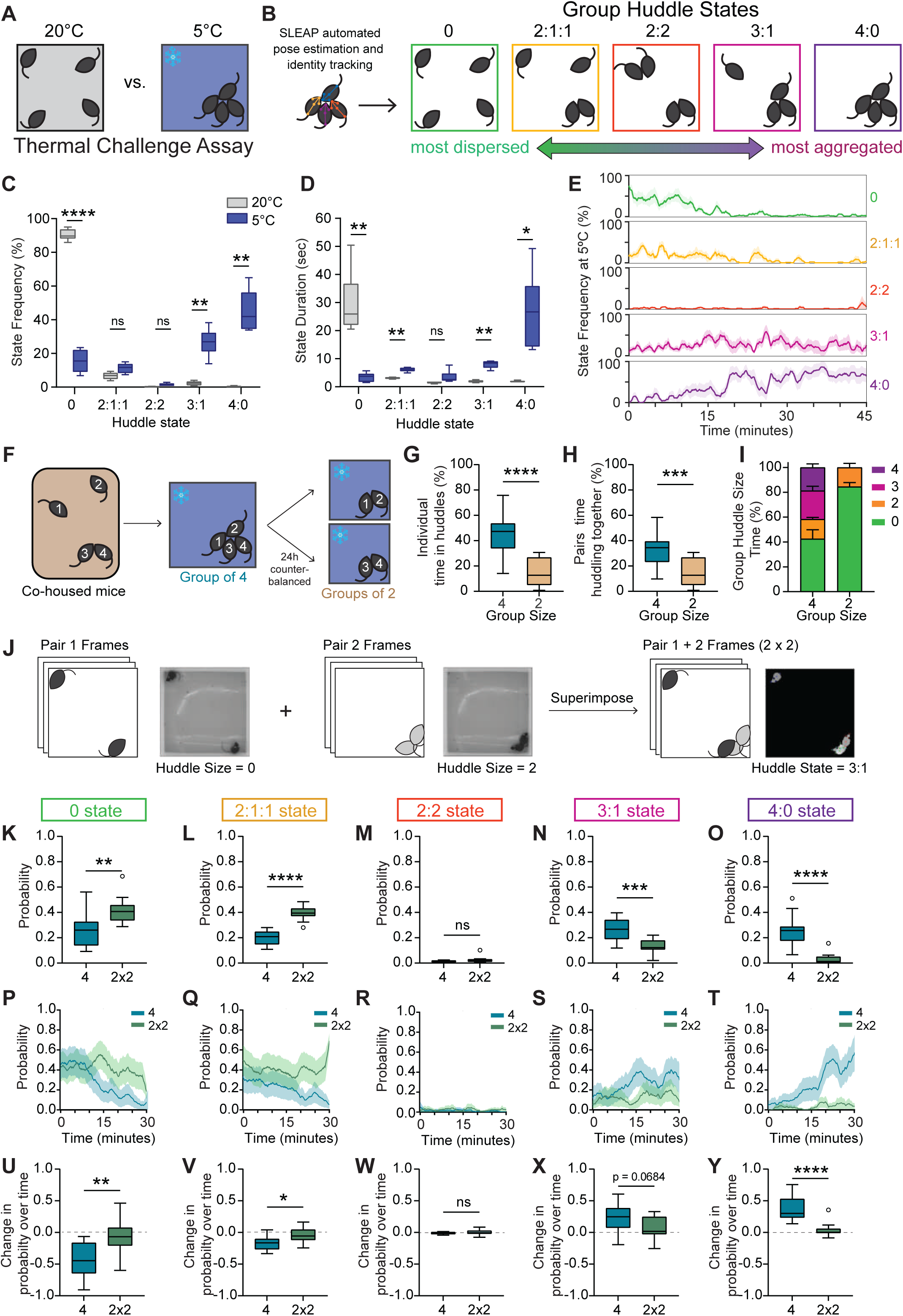
Groups of mice self-organize into huddles in response to cold ambient temperature. **A.** Schematic illustrating groups of four co-housed mice tested in a 40×40cm arena during thermal challenge assay. **B.** Schematics illustrating 5 unique group states derived via automated SLEAP pose estimation and identity tracking, ranging from most dispersed to most aggregated. **C.** Frequency of group states observed at 20°C or 5°C during thermal challenge assay (n = 6 groups of 4 individuals). **D.** Mean group state duration in seconds observed at 20°C or 5°C during thermal challenge assay (n = 6 groups of 4 individuals). **E.** Moving average (mean ± SEM) of percent time of all five group states plotted over time at 5°C (n = 6 groups of 4 individuals). **F.** Schematic illustrating behavioral tests to assess how huddling behavior varies as a function of group size. Co-housed mice are tested at 5°C as a group of four or split into two pairs. Identity of each individual mouse is tracked across both sessions. **G.** Individual animal’s total percent time spent in huddles in groups of four vs groups of two (n = 24 individuals from 6 groups). **H.** Pairs total percent time huddling together in groups of four versus groups of two (n = 12 pairs). **I.** Total percent time (mean ± SEM) observed for each group huddle size (0,2,3,4) in groups of two versus groups of four (n = 6 groups of 4 individuals). **J.** Schematic illustrating image processing pipeline for superimposition of pair videos. Raw video frames from pair 1 are superimposed onto raw video frames from pair 2 to create an artificial group of 4 (2×2). **K-O.** Total probability of huddle states observed for all five group states in real groups (4) versus superimposed groups (2×2) (n = 12 per condition). **P-T.** Moving average (mean ± SEM) plotted over time of probability of huddle states observed in real groups (4) versus artificially superimposed groups (2×2) for all five group states (n = 12 per condition). **U-Y.** Change in probability over time (last five minutes minus first five minutes) for all five group states for real groups (4) versus artificially superimposed groups (2×2) (n = 12 per condition). Box and whisker plots indicate the following: center line – median; box limits – upper and lower quartiles; whiskers – minimum and maximum values. Statistical tests include two-way repeated measures analysis of variance (ANOVA) with Bonferroni post-hoc tests (**C-D**), Wilcoxon matched pairs tests (**G-H**), and Mann-Whitney tests (**K-O**,**U-Y**). \**P*<.05, \*\**P*<.01, \*\*\**P*<.001, \*\*\*\**P*<.0001. See Supplementary Table 1 for details of statistical analyses.

We also tested groups of four co-housed females in the thermal challenge assay. Interestingly, we observed very little huddling in females (Fig. S3A-C). Even at 5°C, females spend most of the time in the 0 state. When compared to males, we observe a statistically significant sex difference in group states at 5°C (Two-way ANOVA Interaction p <.0001, see Table S1). Notably, we did not observe any sex differences in general thermotaxis behavior—males and females exhibit similar amounts of time on a warm corner when placed in a cold arena (Fig. S3D-E). This suggests that the lack of huddling observed in females is not simply due to physiological differences in drive to seek heat. Altogether these data point towards a striking sex difference in collective behavior dynamics which may reflect differing social dynamics or strategies in female versus male groups.

Next, we asked how huddling behavior varies as a function of the size of the group. Do animals huddle more in groups of four, or will they huddle for an equal amount of time even when they are in a smaller group of two? To test this, we tested groups of four co-housed males in the thermal challenge assay at 5°C. We tested the whole group of four all together, or split them into two groups of two (tested separately) in a counterbalanced fashion (Fig. 1F). Remarkably, we found that group size plays a substantial role in modulating huddling behavior. Individual animals spend more than twice as much time huddling when they are in a group of four, as opposed to when they are in a group of two (Fig. 1G). Similarly, when we examine the behavior of specific pairs, we find that a given pair will spend more time huddling together when in a group of four, than when tested in a group of two (Fig. 1H). We then examined huddle size at the level of the whole group and found that the increase in huddling observed in the group-of-four condition is mainly driven by huddles of 3 and 4 animals, not by an increase in time spent in huddles of two (Fig. 1I). These data suggest that group environments create emergent dynamics and behaviors that are not present in pair level interactions.

Next, we wanted to understand whether the increased huddling observed in a group context is simply an artifact of higher density – do animals huddle more in groups of four simply because there are more of them in the same space and interactions are more likely to occur? To test this, we generated videos of artificial groups of four animals (termed ‘2×2’) by superimposing raw frames from videos where only two animals were tested (Methods, Fig. 1J). We then assessed their group huddle states using the approaches described above. Interestingly, we found that real groups of four huddle much more than superimposed 2×2 groups. Groups of four spend less time in the 0 and 2:1:1 states than superimposed 2×2 groups, and more time in the 3:1 and 4:0 states (Fig. 1K-O). We found that for real groups of four, the probability of the 0 state (Fig. 1P,U) and 2:1:1 state (Fig. 1Q,V) decrease throughout the session, while the probability of the 4:0 state significantly decreases (Fig. 1T,Y), and the probability of the 3:1 state trends toward a decrease (Fig. 1S,X). In contrast, state probabilities for superimposed 2×2 groups remain stable throughout the session (Fig. 1P-Y). Together, these data suggest that increased huddling in groups occurs because of emergent social dynamics, and not simply due to a higher likelihood of interaction.

### Individual animals display active and passive decisions to enter and exit huddles

Having characterized the dynamics of collective huddling behavior during the thermal challenge assay, we next wanted to understand the decision-making processes that enable huddles to occur. What types of decisions do individuals make to either engage or disengage in a group huddle? We observed four distinct types of decisions made by individuals to enter and exit huddles, which we characterized into active (self-initiated) and passive (partner-initiated) categories (Fig. 2A). We developed a custom behavior annotation and analysis software, BehaviorAnnotator (https://pypi.org/project/bannotator/), which enables manual experimenter annotation of multi-animal behavior videos (Fig. S4). We annotated active entry decisions as those in which the subject animal approaches another animal(s) and initiates the formation of the huddle. In contrast, passive entry decisions are those in which the subject animal remains stationary and allows another animal to form the huddle. Active exiting decisions are those in which the subject animal leaves the huddle, while passive exiting decisions are those in which the subject animal stays committed to its location, and a partner animal leaves. To further characterize these active and passive behavioral decisions, we used pose estimation data from SLEAP tracking to examine animals’ speed as they make these decisions. We found that during active entry decisions, an individual’s speed is high before huddle onset and rapidly decreases when the animal enters the huddle, while speed is consistently low during passive entry decisions (Fig. 2D). In contrast, during active exiting decisions, an animal’s speed rapidly increases as it exits the huddle, while speed is stably low during passive exiting (Fig. 2G).

**Figure 2:**
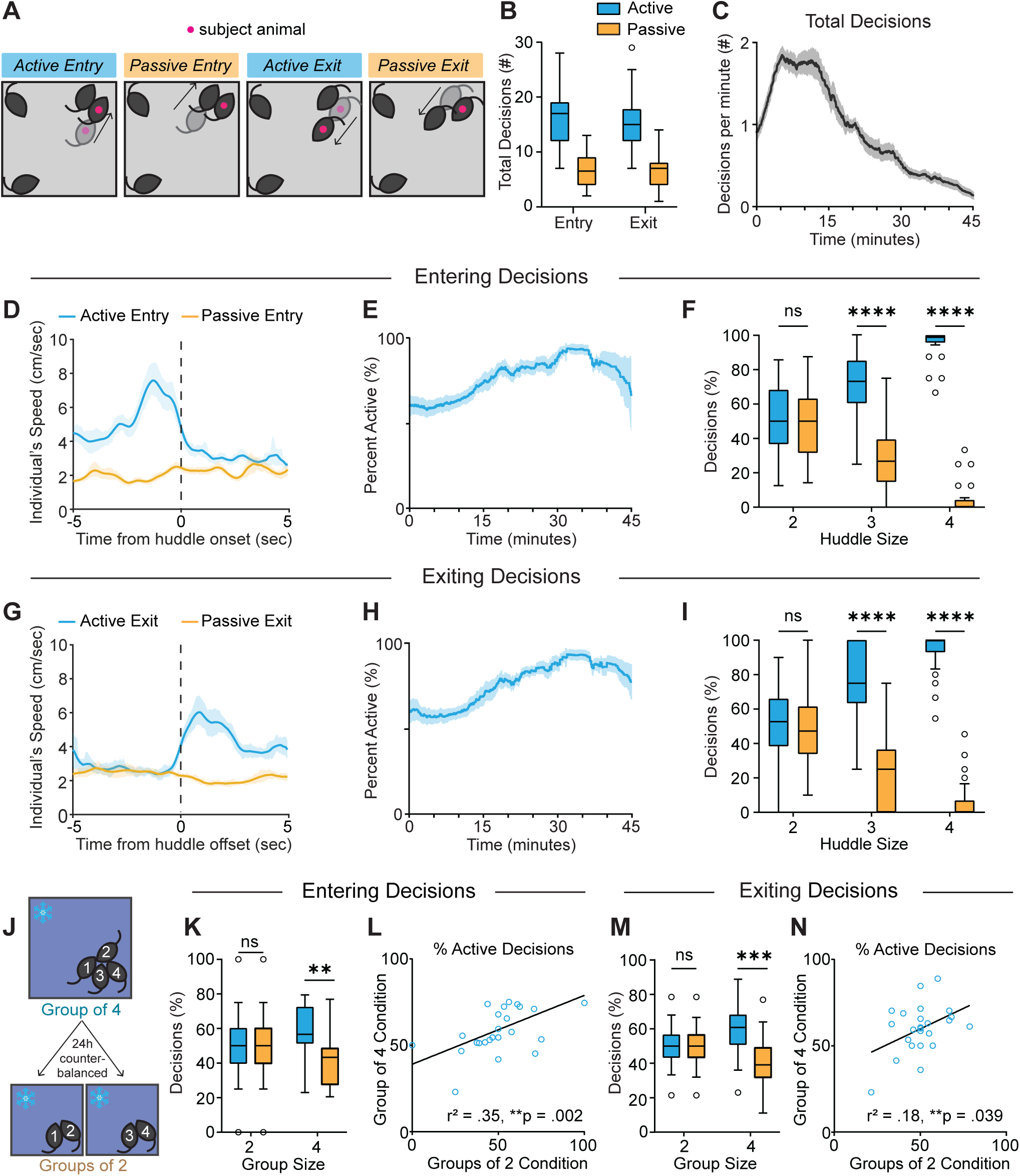
Individual animals display active and passive decisions to enter and exit huddles. **A.** Schematics illustrating active and passive decisions to enter and exit the huddle. Active decisions are self-initiated by the subject animal, while passive decisions are initiated by a partner animal. **B.** Number of decisions observed for all four decision types throughout a session (n = 24 individuals from 6 groups). **C.** Moving average (mean ± SEM) of all four decision types plotted over time for all individuals, smoothed by a .5 sec window (n = 24 individuals from 6 groups). **D,G.** Time course of individual’s speed in (cm/sec) for active and passive entry and exit, respectively, centered to huddle onset or offset (mean ± SEM, n = 11 animals). **E,H.** Moving average (mean ± SEM) of percentage of entry or exit decisions, respectively, that are active plotted over time for all individuals, smoothed by a .5 sec window (n = 24 individuals from 6 groups). **F,I.** Percentage of entry or exit decisions, respectively, that are active versus passive as a function of huddle size (n = 24 individuals from 6 groups). **J.** Schematic illustrating behavioral tests to assess how huddling behavior varies as a function of group size. **K,M.** Percentage of entry or exit decisions, respectively, that are active versus passive as a function of group size (n = 24 individuals from 6 groups). **L, N.** Correlation between percentage of entry or exit decisions that are active during group of two condition vs group of four condition (n = 24 individuals from 6 groups). Box and whisker plots indicate the following: center line – median; box limits – upper and lower quartiles; whiskers – minimum and maximum values. Statistical tests include two-way repeated measures analysis of variance (ANOVA) with Bonferroni post-hoc tests (**F**,**I**,**K**,**M**) and linear regressions (**L**,**N**). \**P*<.05, \*\**P*<.01, \*\*\**P*<.001, \*\*\*\**P*<.0001. See Supplementary Table 1 for details of statistical analyses.

We examined the behavior of 24 unique individuals from 6 groups and found that overall, most of the entry and exiting decisions are active (Fig. 2B). We do observe individual difference in active vs. passive decisions within each group, with some animals showing a higher percentage of active decisions than others (Fig. S5 A-B). Interestingly, we observe a strong correlation between the percentage of active entry decisions and percentage of active exit decisions, suggesting that some animals are stronger initiators of huddle formation and dissolution (Fig. S5C).

When observing the dynamics of decisions across time, we find that the number of decisions made per minute rapidly increases in the first few minutes of the session, and then slowly decreases thereafter (Fig. 2C) as huddles become larger and more stable, and last longer (see Fig. 1E). However, although the number of decisions decreases as the session progresses, the percentage of decisions that are active increases over time (Fig. 2E,H and Fig. S5D,E). We found that this increase in active decisions over time is mainly because passive decisions happen more easily for smaller huddles of two where only one other animal needs to initiate the behavior. The larger a huddle is, the lower the probability of a passive decision, as several other animals must decide to enter or exit. Therefore, if an animal wants to join or exit a larger huddle of 3 or 4, they often must do so actively. Huddles of two are most frequent in early timepoints in the session while huddles of three and four are most frequent later in the session (see Fig. 1E). Accordingly, we found that most passive decisions occur in smaller huddles of two, while huddles of three and four tend to have more active decisions (Fig. 2F,I).

Next, we investigated how decisions that individuals are modulated by group size. Using the same paradigm used to assay the effects of group size described above (Fig. 2J), we found that in groups of two, entry and exiting decisions are roughly half active and half passive – this is mainly because the active decision for one animal is often met with a passive decision for a partner animal (Fig. 2K,M). On the other hand, in a group of four, the majority of entry and exiting decisions are active, because as huddles become larger, a greater number of animals have to simultaneously coordinate for a subject to passively enter or exit (Fig. 2K,M). We found that the percentage of entry decisions that are active in a group of two are strongly correlated with those made in a group of four, suggesting that individuals have stable roles they play in huddle formation in different group conditions (Fig. 2L). In contrast, we observed a weaker correlation in percentage of exiting decisions that were active (Fig. 2N), suggesting that the roles animals play in huddle dissolution are less stable.

### Neural encoding of huddling behaviors

To examine the neural representations of group huddling dynamics, we carried out *in vivo* microendoscopic calcium imaging in the dmPFC of freely behaving animals. Work from our lab and others points towards the prefrontal cortex in encoding one’s own social decisions and the decisions of social partners (40–46). Moreover, accumulating evidence suggests that prefrontal subregions play an important role in encoding group-level social interactions and coordinating one’s behavior in the context of the group (14,41,47,48). However, whether and how dmPFC may encode and regulate group dynamics during huddling remains unclear. Specifically, how neural activity dynamics in the dmPFC may uniquely encode huddling, as well as the active and passive decisions that enable huddles to form, remains elusive. To address these questions, we used calcium imaging to optically record from one animal while it undergoes the thermal challenge assay with its cage mates. We injected an adeno-associated virus (AAV) expressing the fluorescent calcium indicator GCaMP6f and implanted a gradient refractive index (GRIN) lens above the dmPFC (Fig. 3A-C). We recorded a total of 5,141 dmPFC neurons from 11 different animals.

**Figure 3:**
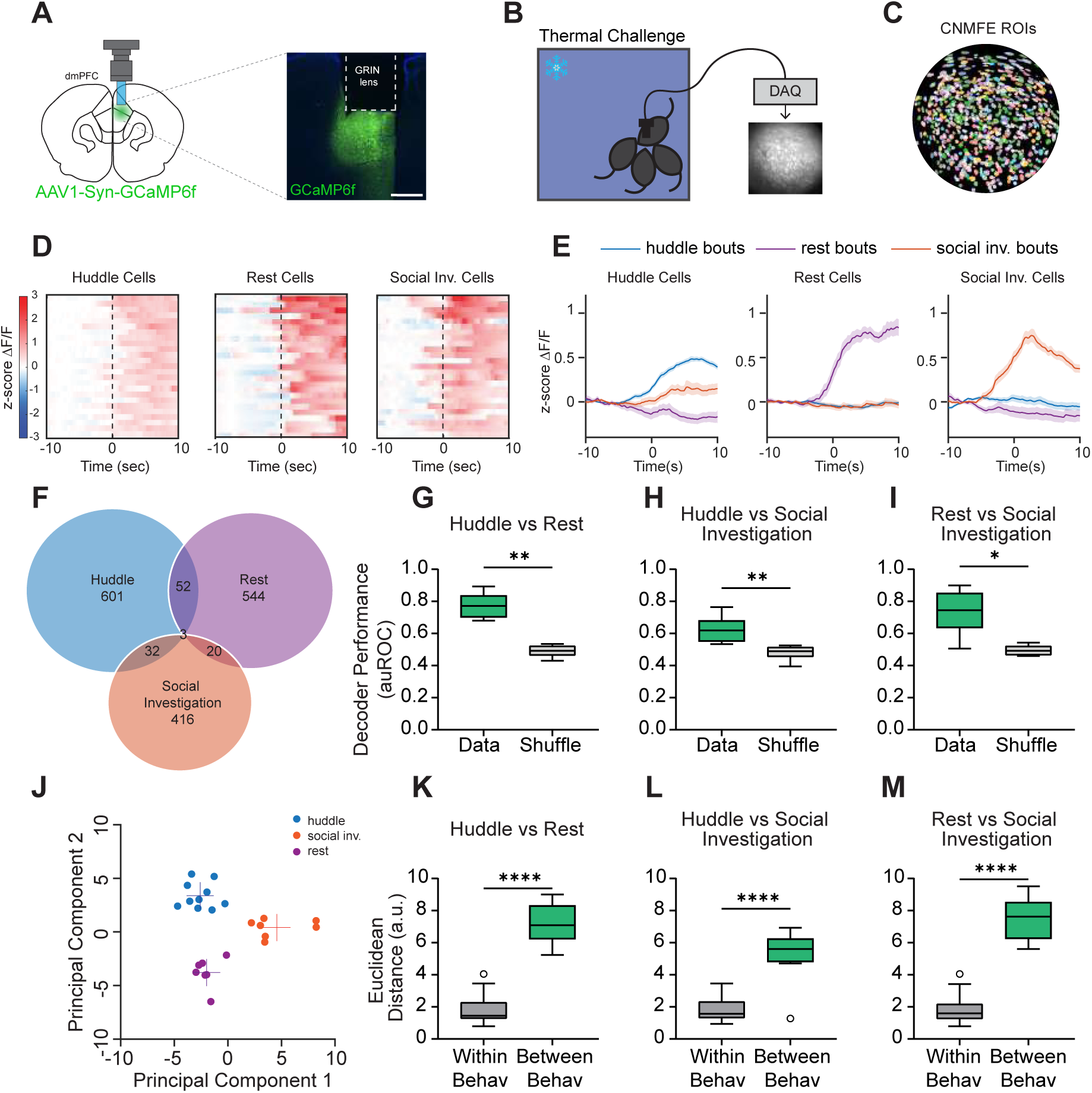
Single cell and population responses of dmPFC neurons during huddling. **A.** Schematic illustrating microendoscopic calcium imaging (left). Example image showing GCaMP6f expression and gradient refractive index (lens) implantation in the dmPFC (right). Scale bar, 500 µm**. B.** Schematic illustrating calcium imaging from one animal and representative imaging field of view during thermal challenge assay. **C.** Single neurons extracted from field of view using CNMF-E. **D.** Heatmaps showing average responses of example huddle cells, rest cells, and social investigation cells activated selectively by their respective behavior, aligned to behavior onset (time 0). **E.** Average responses (mean ± s.e.m.) of cells activated by huddle, rest, and social investigation during huddle bouts, rest bouts, and social investigation bouts. **F.** Venn diagram showing overlap of neurons responsive to huddle, rest, and social investigation. **G-I.** Performance of SVM decoders trained to classify huddling versus rest, huddling versus social investigation, and rest versus social investigation. **J.** Principal component (PC) separation of behavior-evoked population responses from one example imaging session; each dot is the mean response from one behavior bout. a.u., arbitrary units. **K-M.** Average Euclidean distances between PC-projected population vectors within or between huddle and rest behaviors, huddle and social investigation behaviors, or rest and social investigation behaviors. Total # of imaged cells = 5141 from 11 animals. Box and whisker plots indicate the following: center line – median; box limits – upper and lower quartiles; whiskers – minimum and maximum values. Statistical tests include Wilcoxon matched pairs tests (**G-I**), and Mann-Whitney tests (**K-M**). \**P*<.05, \*\**P*<.01, \*\*\**P*<.001, \*\*\*\**P*<.0001. See Supplementary Table 1 for details of statistical analyses.

We first aimed to understand whether dmPFC populations represent the huddling status of the animal, and if so, how this neural representation may differ from that of other similar behavior states. To do this, we compared huddling bouts to bouts of generic non-huddling social investigation (sniffing another animal’s nose, body, or anogenital region) to determine whether neural representations of huddling are different from general social interactions. As an additional control, we also compared them to rest, when the animal is stationary for a prolonged period, since huddling also involves immobility. We then used receiver operating characteristic (ROC) analysis to identify sub-populations of dmPFC neurons that are activated or suppressed by huddling, social investigation, and rest. We found that encoding of these three behaviors is largely separable within the dmPFC. Single neurons in the dmPFC formed largely non-overlapping subpopulations (Fig. 3F) that responded specifically to huddle, social investigation, or rest (Fig. 3D-E), suggesting that huddling behavior may elicit unique population-level neural signatures. To test this, we trained support vector machine (SVM) decoders to classify huddling, rest, and social investigation bouts from each other using dmPFC population activity. We found that the cross-validated performance of these decoders was significantly higher than chance levels (Fig. 3G-I), indicating that unique and stable response patterns in the dmPFC encode huddling behavior. To further explore whether huddling is separable from rest or social investigation, we projected population activity during these behaviors onto principal components; this revealed a clear separation of activity clusters based on behavior type (Fig. 3J). Moreover, the distance between different behaviors was significantly larger than within-behavior distances (Fig. 3K-M), indicating that the separation of responses is not simply due to trial-by-trial variability. Together, these data suggest that the dmPFC contains neural representations of huddling that are distinct from other behavior types.

As a control, we tested whether cold ambient temperature alone non-specifically alters dmPFC encoding of social stimuli by performing microendoscopic calcium imaging of dmPFC neurons in the pencil cup social preference assay at both room temperature and 5°C (Fig. S6A). We did not observe any effect of cold temperature on social preference – animals strongly prefer to interact with a cup containing a social stimulus over a cup containing an inanimate toy regardless of ambient temperature (Fig. S6B). Moreover, we found that dmFPC encodes social and non-social stimuli in distinct neural ensembles at both room temperature and 5°C (Fig. S6C-F). Similarly, population level SVM decoders trained to decode social investigation from toy investigation performed significantly higher than shuffle controls at both room temperature and 5°C (Fig. S6G-H). These data suggest that cold ambient temperature does not generally alter social exploration or dmPFC representations of social stimuli.

### Neural encoding of active and passive decisions

Next, we wanted to understand how dmPFC encodes the active and passive decisions that enable huddles to form and dissolve. To do this, we used ROC analysis to identify single-cell responses to active and passive decisions to enter and exit huddles. Because active entry and active exit decisions necessarily involve dynamic changes in speed as the animal adjusts its behavior state, we used pose estimation data from SLEAP to identify an equivalent number of speed-matched running bouts for both active entry and active exit, for each animal we imaged (Fig. 4B,F). We found that single neurons in dmPFC formed largely non-overlapping subpopulations that responded specifically to active entry, passive entry, and speed-matched running (Fig. 4A,C,D). Similarly, we found that dmPFC neurons encoding active exit, passive exit, and speed-matched running showed little overlap and respond specifically to each behavior (Fig. 4E,G,H). Moreover, the majority of single cells responsive to active and passive decisions do not overlap with other behaviors such as huddle, rest, and social investigation (Fig. S7).

**Figure 4:**
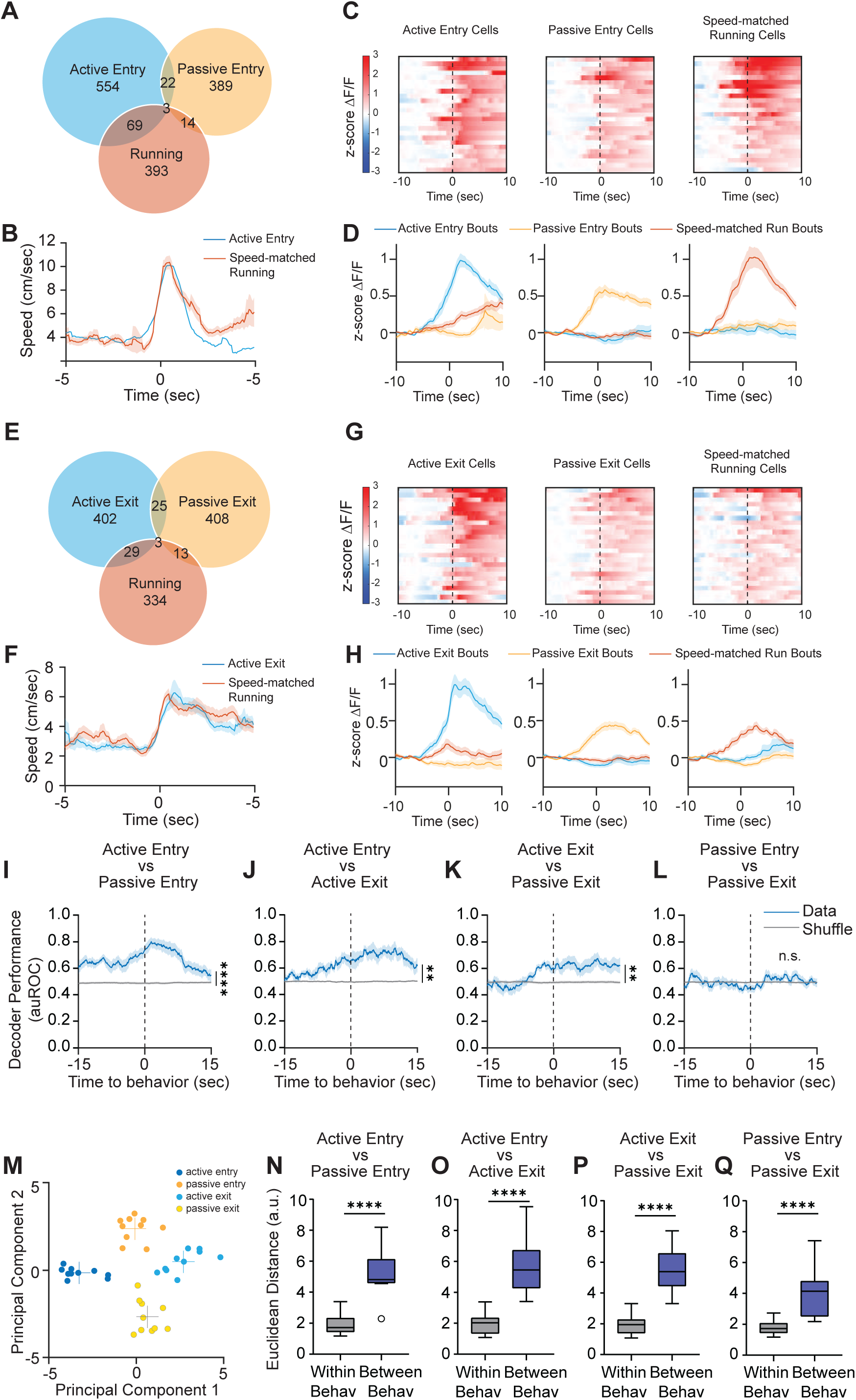
dmPFC encodes active and passive strategies to enter and exit huddles. **A.** Venn diagram showing overlap of neurons responsive to active entry, speed-matched running, and passive entry. **B.** Average speed (mean ± s.e.m.) of subject animal during active entry bouts and speed-matched running bouts identified using SLEAP pose tracking. **C.** Heatmaps showing average responses of example active entry cells, running cells, and passive entry cells activated selectively by their respective behavior, aligned to behavior onset (time 0). **D.** Average responses (mean ± s.e.m.) of cells activated by active entry, running, and passive entry during active entry bouts, running bouts, and passive entry bouts. **E.** Venn diagram showing overlap of neurons to active exit, speed-matched running, and passive exit. **F.** Average speed (mean ± s.e.m.) of subject animal during active exit bouts and speed-matched running bouts identified using SLEAP pose tracking. **G.** Heatmaps showing average responses of example active exit cells, running cells, and passive exit cells activated selectively by their respective behavior, aligned to behavior onset (time 0). **H.** Average responses (mean ± s.e.m.) of cells activated by active exit, running, and passive exit during active exit bouts, running bouts, and passive exit bouts. **I-L.** Performance of SVM decoders trained to classify active entry versus passive entry (**I**), active entry versus active exit (**J**), active exit versus passive exit (**K**), and passive entry versus passive exit (**L**). **M.** Principal component (PC) separation of behavior-evoked population responses from one example imaging session; each dot is the mean response from one behavior bout. a.u., arbitrary units. **N-Q.** Average Euclidean distances between PC-projected population vectors within or between active and passive decisions. Total # of imaged cells = 5141 from 11 animals. Box and whisker plots indicate the following: center line – median; box limits – upper and lower quartiles; whiskers – minimum and maximum values. Statistical tests include two-way repeated measures analysis of variance (ANOVA) (**I-L**) and Mann-Whitney tests (**N-Q**). \**P*<.05, \*\**P*<.01, \*\*\**P*<.001, \*\*\*\**P*<.0001. See Supplementary Table 1 for details of statistical analyses.

Next, we used SVM decoders to examine how separable active and passive decisions are from each other within dmPFC population activity. We constructed SVM decoders that are trained to classify active entry from passive entry (Fig. 4I), active entry from active exit (Fig. 4J), active exit from passive exit (Fig. 4K), and passive entry from passive exit (Fig. 4L). The only decoder which performed at chance was the one trained to decode passive entry from passive exit (Fig. 4L), which may be because in both cases the change in huddle status is driven by the behavior of another animal. All three other decoders performed significantly higher than chance (Fig. 4I-L), and in fact in all cases, decoder performance increases seconds before the onset of the behavior. Similarly, when population activity during these behaviors is projected onto principal components, we observed a clear separation of activity clusters based on behavior type, and the distance between behaviors was greater than the distance within behaviors (Fig. 4M-Q). Because decoding active and passive decisions from each other can simply reflect locomotor artifacts, we also trained SVM decoders to classify active decisions from running and passive decisions from rest and found that these decoders perform higher than chance (Fig. S8A-D). Collectively, these data suggest that dmPFC population activity can decode active and passive decisions from each other in a manner that is not simply reflective of locomotor activity.

Interestingly, although decoders could effectively predict active decisions from passive decisions, we found that there was some shared encoding of active entry and active exit, as well as passive entry and passive exit (Fig. S8E-L). We trained SVM decoders on one behavior type and tested them on another type to determine how much shared information there is within dmPFC about these decisions. We found that SVMs trained on active entry can predict active exit higher than chance (and vice versa, Fig. S8E-F). Likewise, SVMs trained on passive entry can predict passive exit higher than chance (and vice versa, Fig. S8G-H). However, decoders trained on active decisions are not effective at predicting passive decisions, and decoders trained on passive decisions are not effective at predicting passive decisions (Fig. S8I-L). Together, these data suggest that active decisions generally rely on some partially shared encoding in dmPFC, which is distinct from partially shared encoding between passive decisions.

Interestingly, although we were able to decode many aspects of active and passive decisions, we were not able to decode higher order features of group huddles. Although individual animals demonstrate preferences to huddle with particular other animals in the group (Fig. S9A-D), we were not able to decode information about the membership of the huddle from dmPFC population activity (Fig. S9E-F). Likewise, we were also not able to decode information about the size of the huddle (Fig. S9G-J). Collectively, these data suggest that the main role of dmPFC in coordinating group behavior is to track the decision-making processes of the subject animal as well as social partners.

### Chemogenetic silencing of dmPFC alters active and passive behavior decisions in manipulated subjects as well as non-manipulated partners

The neural representation of huddling behaviors raises the possibility that dmPFC causally controls group huddling in the thermal challenge assay. To test this hypothesis, we virally injected an AAV expressing hM4Di, an inhibitory chemogenetic effector, under a CaMKii promoter to silence dmPFC principal neurons (Fig. 5A). We injected hM4Di bilaterally into dmPFC in all four animals per cage (Fig. 5B). This allowed us to run the same groups on multiple days, alternating administration of clozapine-N-oxide (CNO) drug or saline to achieve within animal control. We tested two experimental conditions: 4SAL in which all four animals receive saline, or 2CNO,2SAL, in which two of the animals receive CNO and the other two receive saline. We counterbalanced the 2CNO,2SAL condition so that all animals receive both saline and CNO (Fig. 5B). This experimental design allowed us to ask two important questions: 1. Compared to baseline, how does silencing dmPFC alter the behavior of the CNO-injected animal during group huddling? and 2. How does silencing dmPFC in some members of the group change the behavior of the other non-manipulated animals?

**Figure 5:**
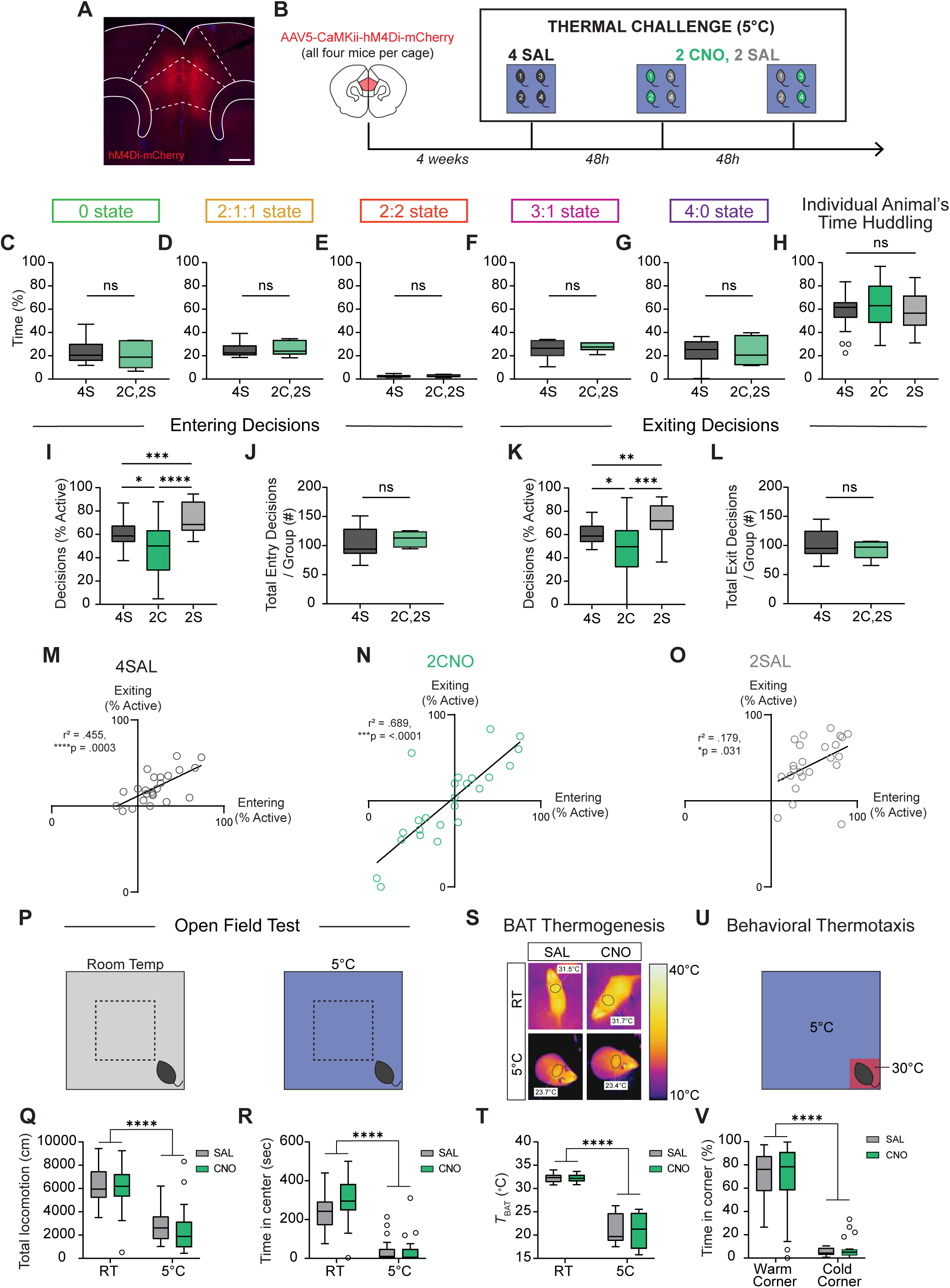
Chemogenetic silencing of dmPFC alters active and passive behavioral decisions in subject animals as well as their partners. **A.** Example image showing AAV-hM4Di-mCherry expression in the dmPFC. Scale bar, 500 µm. **B.** Schematic illustrating experimental paradigm for DREADD inhibition of dmPFC during thermal challenge. 4 SAL refers to condition in which all four animals are injected with saline. 2 CNO, 2 SAL refers to condition in which two animals are injected with CNO, and two with saline. **C-G.** Percent time in huddle states observed for all five group states during 4S and 2C,2S conditions (n = 6 groups). **H.** Individual animal’s total percent time spent huddling in 4S, 2C, and 2S conditions (n = 24 individuals from 6 groups). **I,K.** Within animal comparison of percent of entry or exit decisions, respectively that are active during 4S, 2C, and 2S conditions (n = 24 individuals from 6 groups). **J,L.** Total number of entry or exit decisions, respectively per group during 4S and 2C,2S conditions (n = 24 individuals from 6 groups). **M-O.** Correlation between percent of entry decisions and percent of exiting decisions that are active for 4SAL, 2CNO, and 2SAL conditions (n = 24 individuals from 6 groups). **P.** Schematic illustrating chemogenetic inhibition of dmPFC during open field test at room temperature (RT) and 5°C. **Q-R.** Within animal comparison of total locomotion and time in center, respectively, during open field test at room temperature and 5°C after SAL or CNO injection (n = 24 animals **S.** Representative infrared thermal images demonstrating temperature above BAT (brown adipose tissue, black circles) after SAL or CNO injection. **T.** Quantification of thermography images in regions above BAT after SAL or CNO injection (n = 24 animals at RT, 12 animals at 5°C). **U.** Schematic illustrating chemogenetic inhibition of dmPFC during behavioral thermotaxis assay. **V.** Within animal comparison of percent time spent in warm corner versus the average of three cold corners after SAL or CNO injection (n = 24 animals). Box and whisker plots indicate the following: center line – median; box limits – upper and lower quartiles; whiskers – minimum and maximum values. Statistical tests include one-way (**H**,**I**,**K**) and two-way (**Q**,**R**,**T**,**V**) repeated measures analysis of variance (ANOVA) with Bonferroni post-hoc tests, Wilcoxon matched pairs tests (**C-G**,**J**,**L**), and linear regression tests (**M-O**). \**P*<.05, \*\**P*<.01, \*\*\**P*<.001, \*\*\*\**P*<.0001. See Supplementary Table 1 for details of statistical analyses.

We first asked how chemogenetic silencing of dmPFC affects total huddling behavior. Surprisingly, we found no differences in the overall group huddle states between 4SAL and 2CNO, 2SAL conditions (Fig. 5C-G). We further assessed whether dmPFC silencing affects the membership of huddles in the 2CNO,2SAL condition and found that injecting two animals with CNO did not bias huddles of two towards 2 saline animals, 2 CNO animals, or one of each (Fig. S10A-B). We also observed no difference in individual animal’s huddling time (Fig. 5H).

The lack of chemogenetic effect on overall huddling time led us to next ask whether dmPFC silencing may alter behavior in a more specific way by driving changes in active and passive decisions. Indeed, we found that compared to the baseline (4SAL) condition, the animals injected with CNO (2CNO) showed a significant decrease in the percentage of active entry and active exit decisions (Fig. 5I,K). This suggests that dmPFC activity is necessary for active decision-making during group huddling. Remarkably, however, we also found that dmPFC silencing in two animals in the group also influenced the behavior of non-manipulated partners. We found that the saline-injected social partners of the CNO animals (2SAL) showed an increase of equal magnitude in the percentage of active decisions, despite not receiving direct neural manipulation—that is, while CNO-injected animals show a 25% reduction in active decisions, their saline injected-partners show a corresponding 25% increase (Fig. 5I-K). As such, the total sum of all active and passive entry and exit decisions across the group remains constant in both conditions (Fig. 5J,L). These effects on active decisions were mirrored by an equivalent but opposite effect on passive decisions in all conditions (Fig. S10C-D). For the 2CNO,2SAL condition, we also performed a within-group comparison and found that CNO animals show significantly fewer active decisions than their saline partners (Fig. S10E-F). Like Figure 2, we observed considerable variability in individual’s active entry and exit decisions within each condition (Fig. 5M-O). Yet, within all conditions, there remained a positive correlation between the percentage of active entry decisions and the percentage of active exit decisions (Fig. 5M-O). Together, these data suggest that no changes in overall huddling time are observed because the 2SAL animals compensate for the 2CNO animals’ decrease in active decisions, such that overall huddling time is conserved, preserving the homeostatic state of the group.

We next asked whether changes in active decision-making might simply be explained by non-specific alteration in locomotion or anxiety. To rule out this possibility, we assessed each individual’s speed during null periods in the thermal challenge assay – those periods where huddling, entry, or exit decisions did not occur and the animal was freely ambulating in the arena. We did not observe any generalized differences in locomotion (Fig. S10G). To further validate this finding, we chemogenetically silenced dmPFC in individual animals in the Open Field Test at both room temperature and 5°C (Fig. 5P). While we observed a significant main effect of temperature on total locomotion – animals ambulate much less in cold ambient temperature to conserve energy, there was no effect of dmPFC silencing (Fig. 5Q, Fig. S10H). Similarly, we observed a main effect of temperature on measures of anxiety (time in center), but no effect of CNO injection (Fig. 5R). We also use infrared thermal imaging to measure the temperature above the brown adipose tissue (49) (BAT) at room temperature and 5°C (Fig. 5S). We found that BAT temperature is generally much lower at 5°C due to heat loss (Fig. 5T), as acute cold exposure largely drives shivering thermogenesis, and animals need to be exposed to cold for one week or longer to drive BAT activation (50,51). However, we do not observe any effects of dmPFC silencing (Fig. 5T). Finally, we silenced dmPFC in individual animals in a behavioral thermotaxis assay to determine whether dmPFC silencing alters general warm-seeking behaviors (Fig. 5U). Consistent with previous reports, we found no effect of dmPFC silencing on time spent in the warm corner (Fig. 5V) (52). We also tested control animals expressing AAV-CaMKii-mCherry and observed no effects on active decisions, passive decisions, gross huddling time, locomotion, anxiety, BAT thermogenesis or behavioral thermotaxis (Fig. S11). Altogether, these data suggest that silencing dmPFC in individuals within a social group alters collective decision-making in the thermal challenge assay without altering anxiety, locomotion, autonomic thermogenesis, or warm-seeking behaviors.

## DISCUSSION

Here we investigated the neurobiology underlying collective behavior, using thermoregulatory huddling in rodents as a model behavior. Although huddling behavior has been studied in the wild and in the laboratory in the past, we established a behavioral and quantitative framework for understanding group huddle states and the decision-making processes of each individual animal. We demonstrated that huddling behavior varies as a function of the size of the group and that animals huddle more when in a group of four than in pairs. Moreover, we identified active and passive decision-making processes that individual animals make to enter or exit huddles and characterized the dynamics of these decisions over time and in relation to huddle size. These findings establish a novel and accessible foundation for studying the neurobiology of collective behavior in laboratory settings. Traditional model organisms and behaviors studied in collective behavior are not always amenable to neurobiological recordings and manipulations, either due to difficulty studying the behavior of interest indoors (e.g. stampeding, flocking), or difficulty integrating the behavior of interest with currently available tools for neural recordings in some species (for example, insect swarms can be studied indoors, but neural recordings in insects often require immobilizing the animal in a head-fixed position).

Using microendoscopic calcium imaging, we found that dmPFC contains unique populations of neurons that encode active and passive decisions to enter and exit huddles, consistent with previous reports suggesting that dmPFC encodes the behavior of self and other (42,43). These findings suggest that distinct populations of neurons encode one’s own decisions to enter or exit a huddle, versus the decisions of other members of the group. Importantly, encoding of these behavior variables were not simply reflective of differences in locomotor behavior of the animal.

Using chemogenetic silencing, we silenced dmPFC in two out of four animals in the group and found that dmPFC-silenced animals show a reduction in active decision making. That is, animals become more passive and allow other individuals to initiate the formation and dissolution of the huddle. Remarkably, we found that these manipulations not only change the behavior of the two manipulated animals, but also have a downstream effect on non-manipulated partners, despite not receiving a direct neural perturbation. In response, non-silenced partners adjust their behavior and increase their active decision making to compensate for the dmPFC-silenced animals. What changes when the dmPFC is silenced is who actively initiates huddle formation, rather than affinity to huddle. The increase in active decisions on the part of non-manipulated animals may reflect motivation to huddle due to their own unmet physiological needs, or could reflect prosocial decision making to ensure that the overall thermoregulatory homeostatic needs of the whole group are met. Further studies that compare the thermal needs of the manipulated and non-manipulated animals may clarify these differences. These data are the first, to our knowledge, to show that manipulating neural activity in individual animals changes not only their own behavior, but also the behavior of their social partners. These findings point to an important future research focus to understand the neural mechanisms that enable animals to sense changes in the behavior of other individuals and appropriately compensate in response.

Huddling requires animals to sense the behavior of other animals, determine one’s own homeostatic need, and appropriately adapt their own behavior considering both these factors (7). Future research that examines how neural populations integrate the internal state of the animal with information about other animal’s behaviors to guide decision making in a collective context will be promising. Additionally, huddling in the thermal challenge assay very likely depends on thermosensory inputs that convey information about ambient temperature to dmPFC to guide appropriate decisions. Future studies that examine the function of thermosensory inputs to prefrontal regions may be promising in this regard (25,53). It also remains unknown how dmPFC function may differ from or overlap with the role of subcortical circuits in group huddling, especially hypothalamic nuclei that regulate behavioral and autonomic thermoregulation, such as lateral hypothalamus (24) and medial preoptic area (54). Future studies that examine the roles of these brain regions will help map the distinct contributions of brain-wide circuits in mediating collective huddling. Further, while we observed very little huddling in adult females, existing studies have found substantial amounts of huddling in females at earlier stages of development (29,55). The physiological and circuit mechanisms that guide this switch in behavior over development are unknown. These and other avenues will be promising areas of inquiry for collective neuroscience researchers. Moreover, mice huddle not only for thermoregulatory purposes, but also while sleeping (56) and for protection in the presence of threats or predator cues (57,58). To what extent the behavioral and neural mechanisms are conserved across these contexts are open questions for future studies.

Altogether, our study advances understanding into the neurobiology of collective behavior, and sheds light on how the decision-making processes of individuals are rooted in neural ensembles in the brain. These insights enrich our understanding of how social groups respond to environmental challenges and could lead to developments that enhance collective behavior in response to challenges at the level of human society.

**Supplementary Figure 1:**
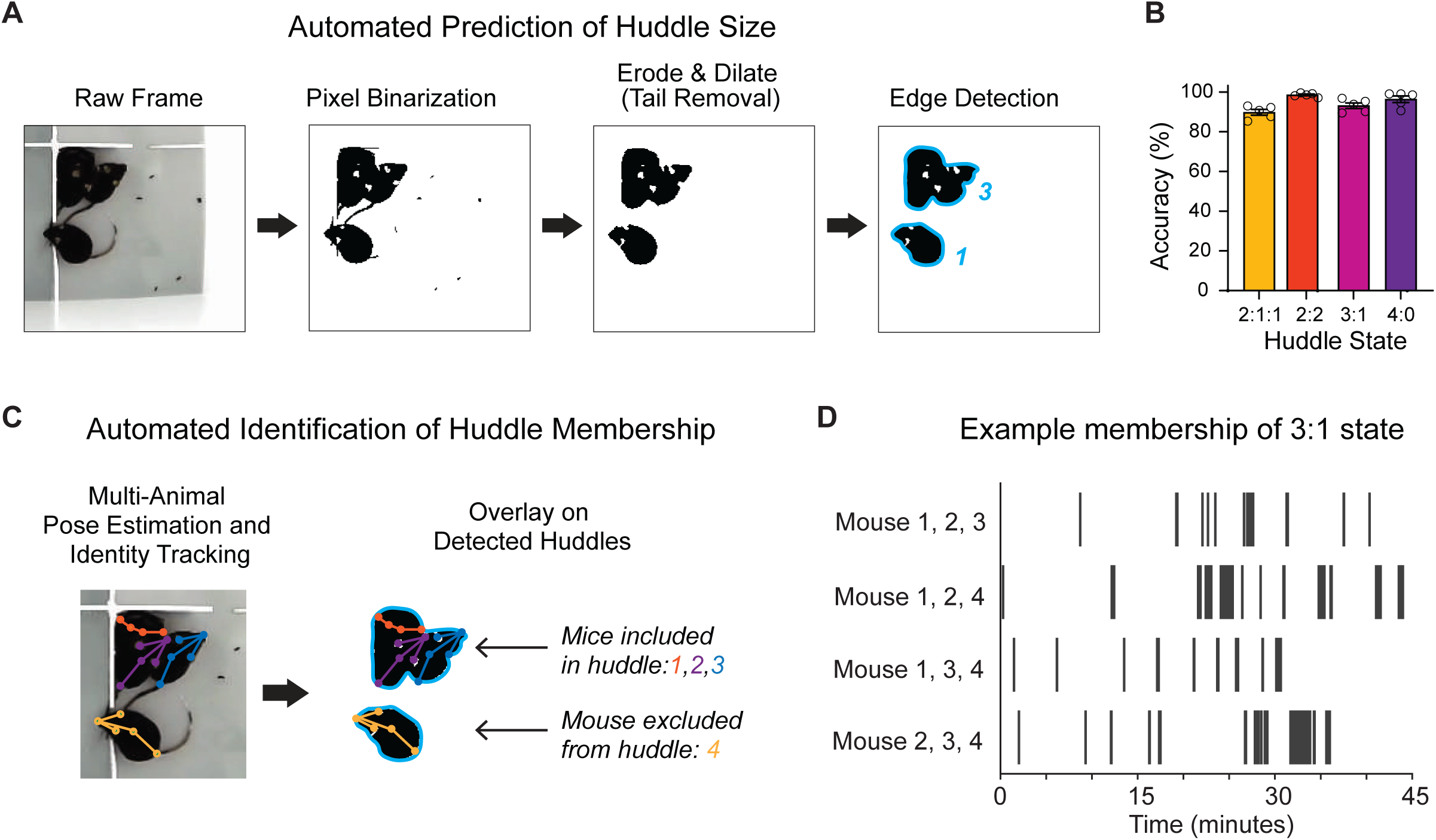
Automated pipeline for group huddle behavior analysis. **A.** Pipeline for automated detection of huddle size. Raw frames are binarized into black and white pixels. Opening (erosion followed by dilation) is performed to removed tails and fecal artifacts. Edge detection is performed to identify connected groups of animals. **B.** Percent accuracy of detected huddle state compared to manual human annotation **C.** Automated Identification of huddle membership is achieved by tracking raw behavior videos with a trained neural network (Social Leap Estimates Animal Poses) to identify individual nodes and identities. Tracked poses and identities are overlayed on top of detected huddles to identify the membership. **D.** Example raster plot for one group demonstrating membership configurations for huddles of three throughout one behavior session.

**Supplementary Figure 2:**
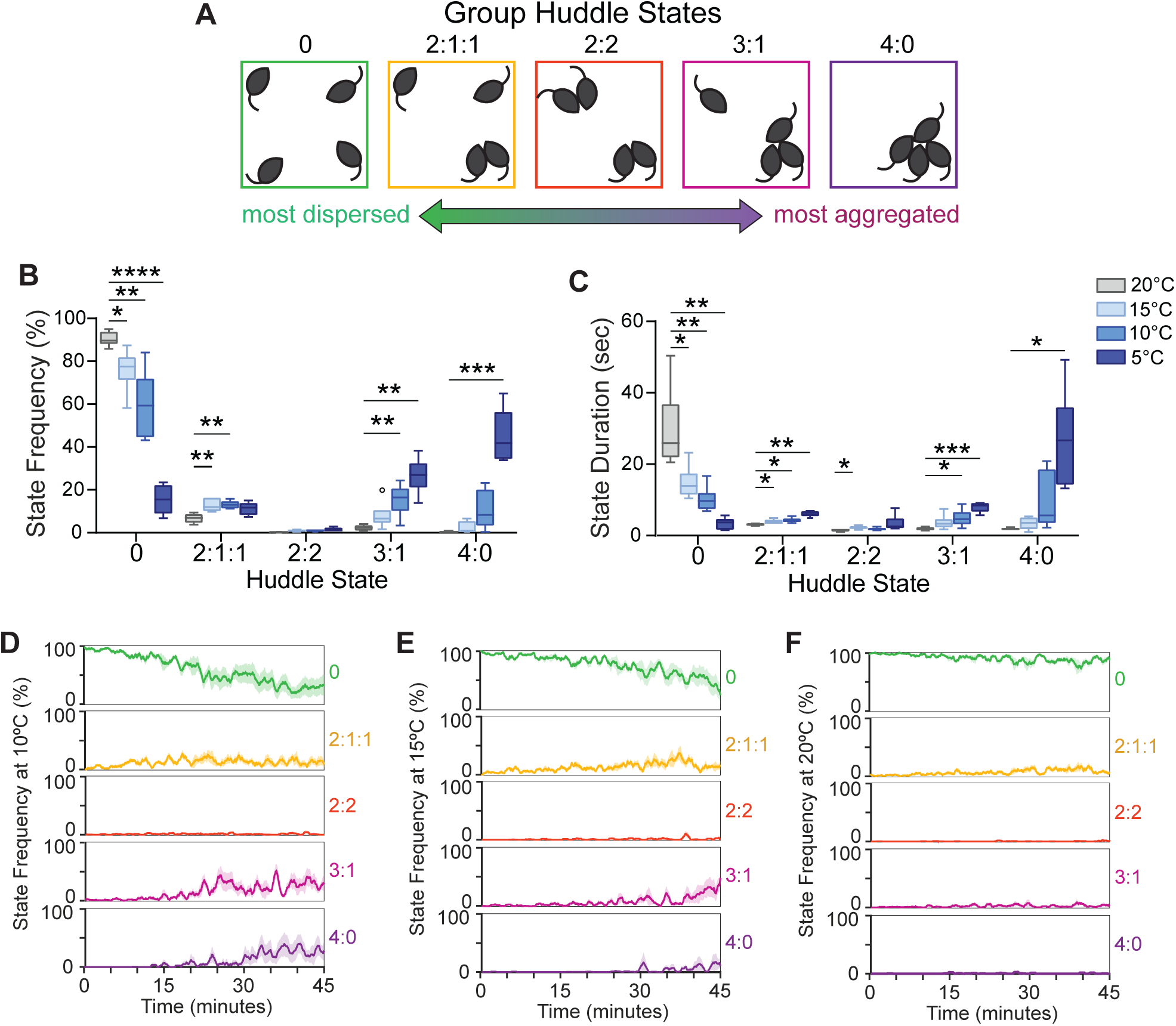
Titration of ambient temperature during thermal challenge assay. **A.** Schematics illustrating 5 unique group states derived via automated SLEAP pose estimation and identity tracking, ranging from most dispersed to most aggregated. **B.** Frequency of group states observed at 20°C, 15°C, 10°C, or 5°C during thermal challenge assay (n = 6 groups of 4 individuals). **C.** Mean group state duration in seconds observed at 20°C, 15°C, 10°C, or 5°C during thermal challenge assay (n = 6 groups of 4 individuals). **D.** Moving average (mean ± SEM) of percent time of all five group states plotted over time at 10°C (n = 6 groups of 4 individuals). **E.** Moving average (mean ± SEM) of percent time of all five group states plotted over time at 15°C (n = 6 groups of 4 individuals). **F.** Moving average (mean ± SEM) of percent time of all five group states plotted over time at 20°C (n = 6 groups of 4 individuals). Box and whisker plots indicate the following: center line – median; box limits – upper and lower quartiles; whiskers – minimum and maximum values. Statistical tests include two-way repeated measures analysis of variance (ANOVA) with Bonferroni post-hoc tests (**B**,**C**). \**P*<.05, \*\**P*<.01, \*\*\**P*<.001, \*\*\*\**P*<.0001. See Supplementary Table 1 for details of statistical analyses.

**Supplementary Figure 3:**
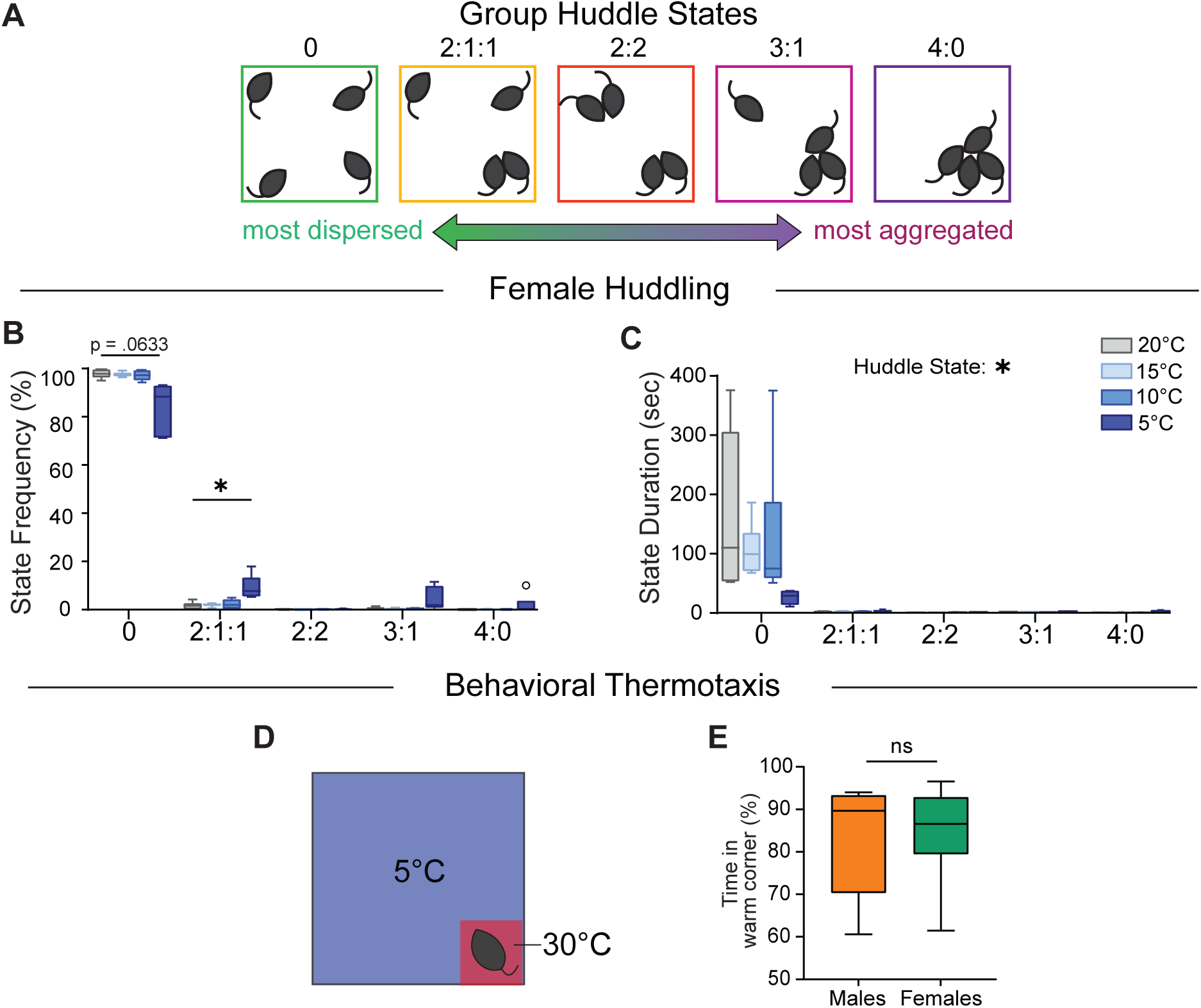
Huddling during thermal challenge in females. **A.** Schematics illustrating 5 unique group states derived via automated SLEAP pose estimation and identity tracking, ranging from most dispersed to most aggregated. **B.** Frequency of group states observed at 20°C, 15°C, 10°C, or 5°C during thermal challenge assay in females (n = 6 groups of 4 individuals). **C.** Mean group state duration in seconds observed at 20°C, 15°C, 10°C, or 5°C during thermal challenge assay in females (n = 6 groups of 4 individuals). **D.** Schematic illustrating behavioral thermotaxis assay. Animals are placed in a behavioral chamber at 5°C with free access to a 30°C warm corner. **E.** Comparison of percent time spent in warm corner during thermotaxis assay in males and females (n = 8 males, 8 females). Box and whisker plots indicate the following: center line – median; box limits – upper and lower quartiles; whiskers – minimum and maximum values. Statistical tests include two-way repeated measures analysis of variance (ANOVA) with Bonferroni post-hoc tests (**B**,**C**), and Wilcoxon matched pairs tests (**E**). \**P*<.05, \*\**P*<.01, \*\*\**P*<.001, \*\*\*\**P*<.0001. See Supplementary Table 1 for details of statistical analyses.

**Supplementary Figure 4:**
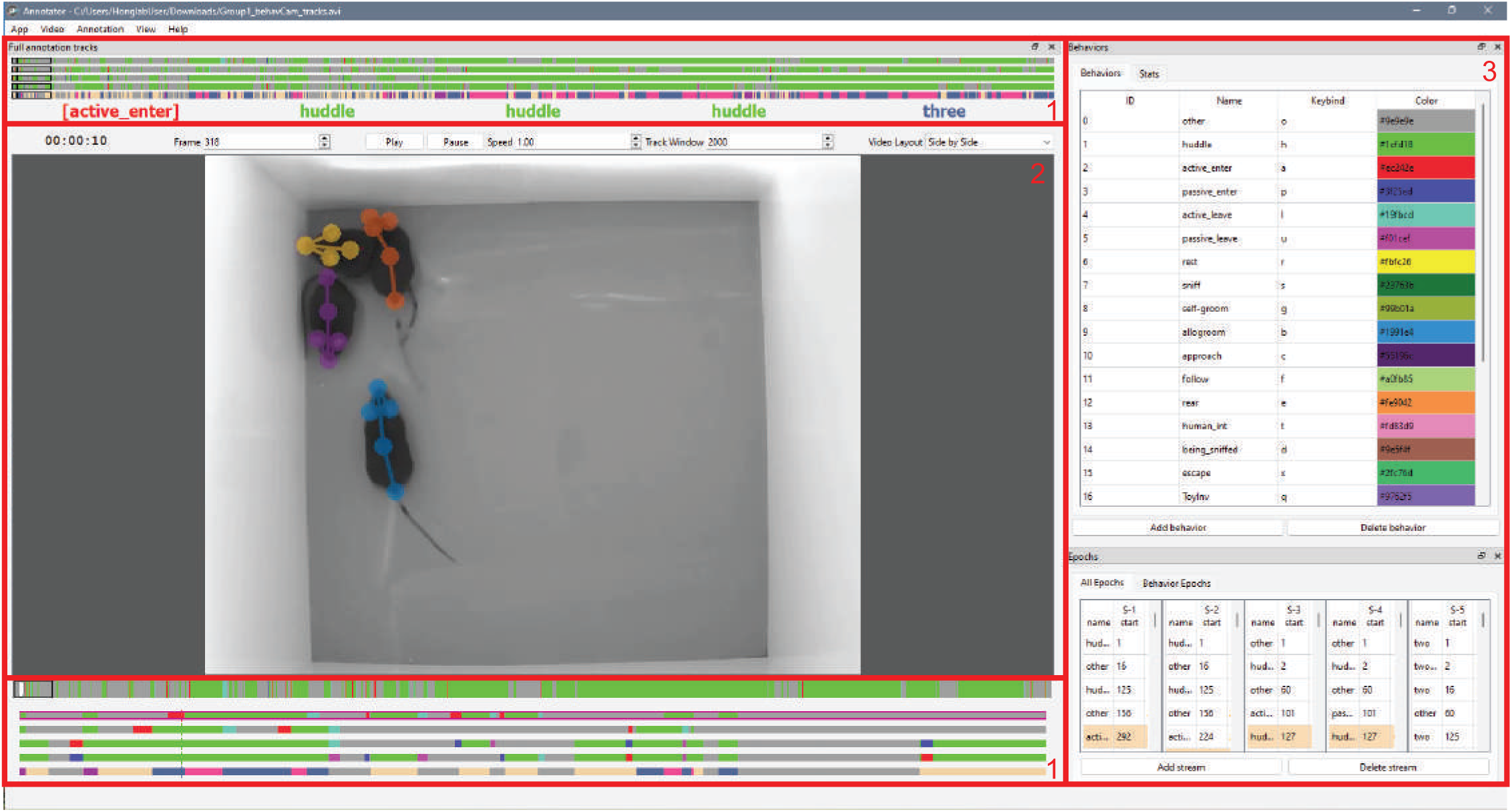
Graphical user interface for BehaviorAnnotator, a custom software for manual annotation and analysis of multi-animal behavior. The graphic user interface of the annotator has 3 panels. Panel 1 displays the annotation streams containing user defined behaviors for all four animals, and a fifth stream which denotes the aggregate huddle size when a huddle is present. Panel 2 displays the behavior video(s). Panel 3 displays the list of user-defined behaviors and labeled behavior epochs.

**Supplementary Figure 5:**
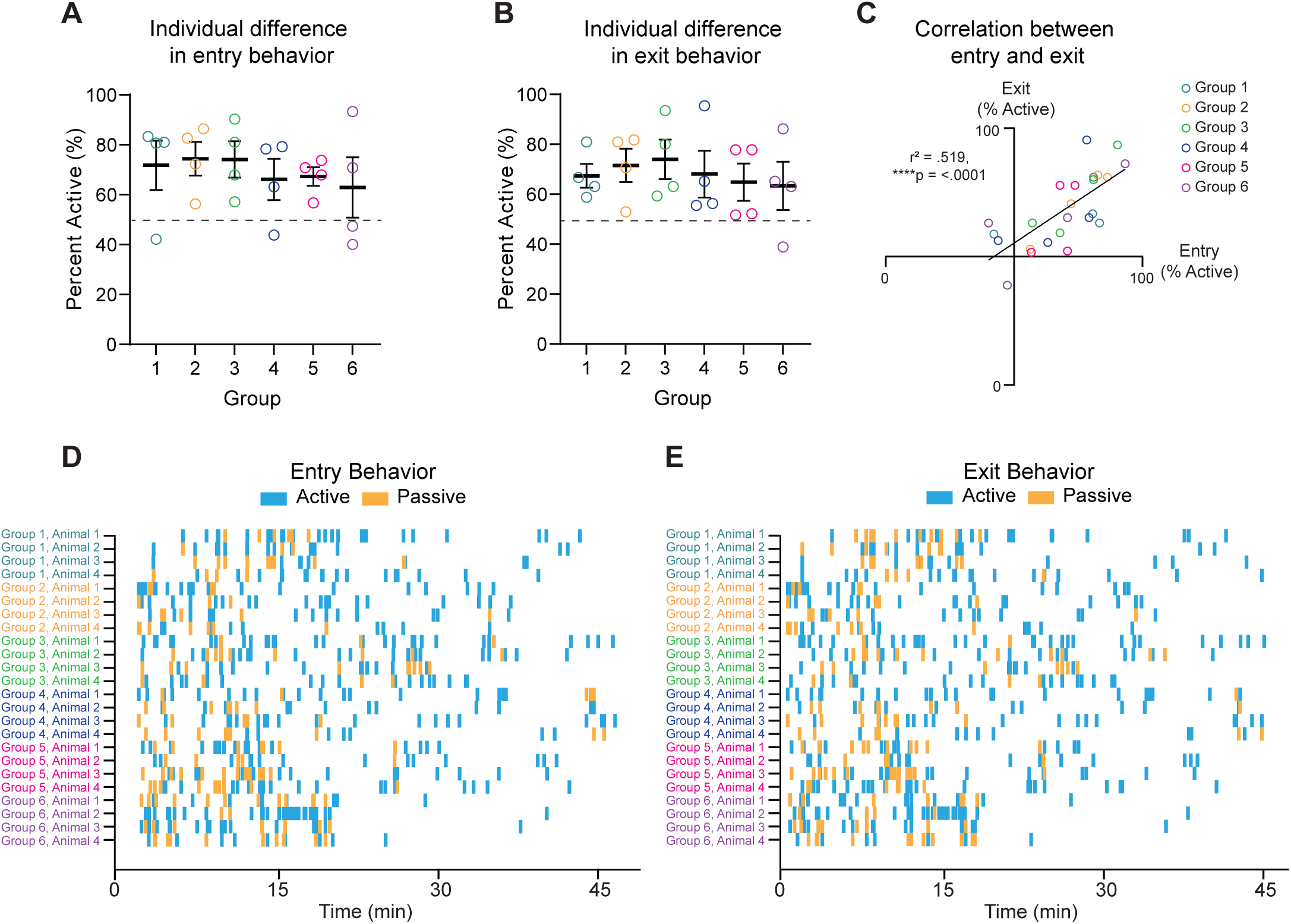
Individual difference in active vs passive decisions. **A.** Percent of entry decisions (mean ± SEM) that are active plotted for all four individuals in six groups. **B.** Percent of exiting decisions (mean ± SEM) that are active plotted for all four individuals in six groups. **C.** Correlation between percent of entry decisions and percent of exiting decisions that are active (n = 24 individuals from 6 groups). **D.** Raster plot illustrating active and passive entry events throughout full behavioral session (n = 24 individuals from 6 groups). **E.** Raster plot illustrating active and passive exiting events throughout full behavioral session (n = 24 individuals from 6 groups). Statistical tests include linear regression (**C**). \**P*<.05, \*\**P*<.01, \*\*\**P*<.001, \*\*\*\**P*<.0001. See Supplementary Table 1 for details of statistical analyses.

**Supplementary Figure 6:**
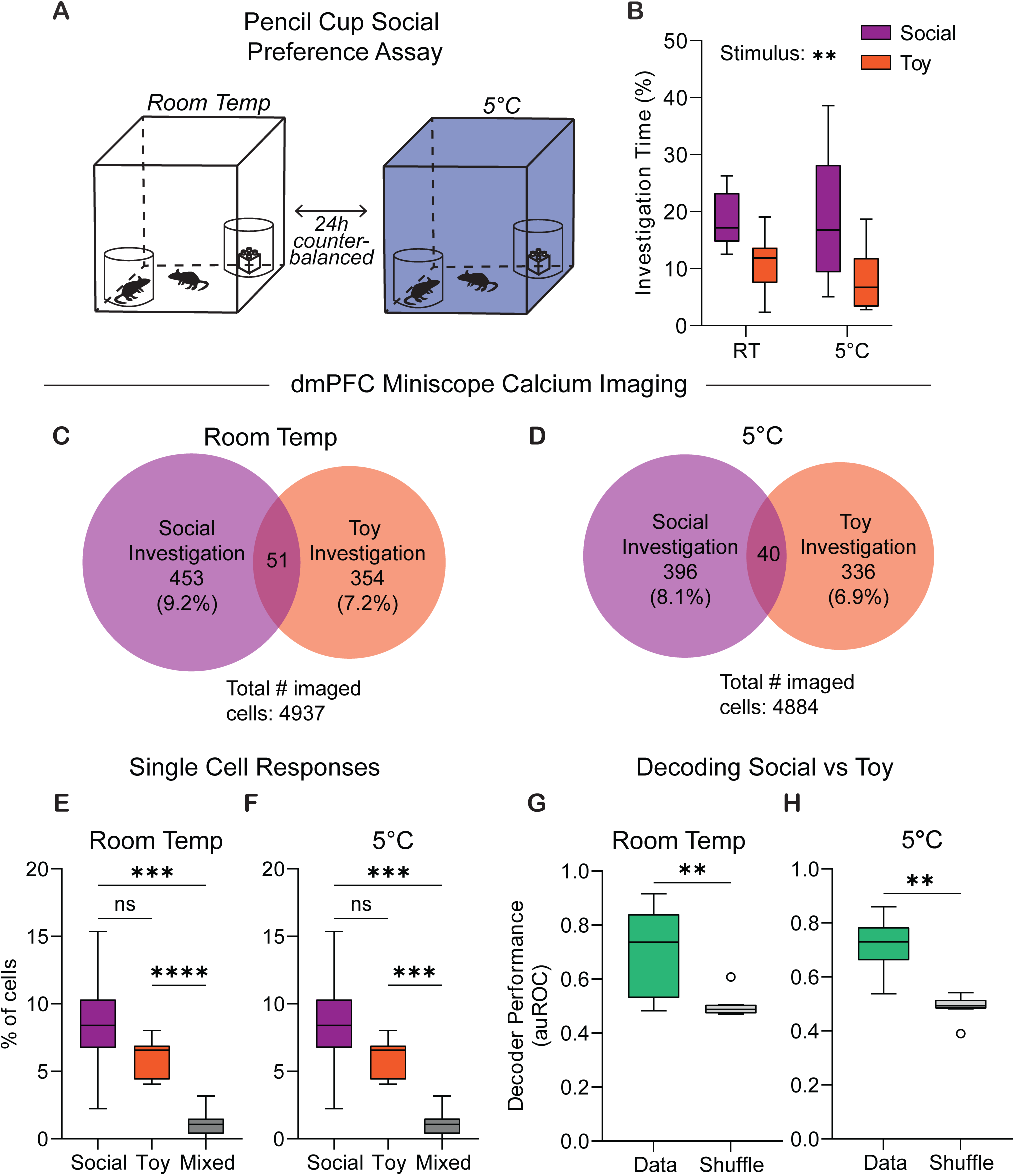
Cold ambient temperature does not alter general social preference or dmPFC encoding of social stimuli. **A.** Schematic illustrating pencil cup social preference assay. Animals were tested for 30 minutes at room temperature or 5°C to determine preference for wired pencil cup containing a conspecific vs a toy. **B.** Quantification of investigation time directed towards social cup vs toy cup at room temperature (RT) or 5°C (n = 10 animals). **C.** Venn diagram showing dmPFC cells responsive to social and toy investigation at room temperature. Total # of imaged cells = 4937 from animals. **D.** Venn diagram showing dmPFC cells responsive to social and toy investigation at room temperature. Total # of imaged cells = 4884 from 10 animals. **E.** Percent of dmPFC cells that are social responsive, toy responsive, or mixed responsive at room temperature (n = 10 animals). **F.** Percent of dmPFC cells that are social responsive, toy responsive, or mixed responsive at 5°C (n = 10 animals). **G.** Support vector machine (SVM) decoder performance to decode social vs toy investigation at room temperature (n = 10 animals). **H.** Support vector machine (SVM) decoder performance to decode social vs toy investigation at 5°C (n = 10 animals). Box and whisker plots indicate the following: center line – median; box limits – upper and lower quartiles; whiskers – minimum and maximum values. Statistical tests include one-way (**E-F**) and two-way (**B**) repeated measures analysis of variance (ANOVA) with Bonferroni post-hoc tests, and Wilcoxon matched pairs tests (**G-H**). \**P*<.05, \*\**P*<.01, \*\*\**P*<.001, \*\*\*\**P*<.0001. See Supplementary Table 1 for details of statistical analyses.

**Supplementary Figure 7:**
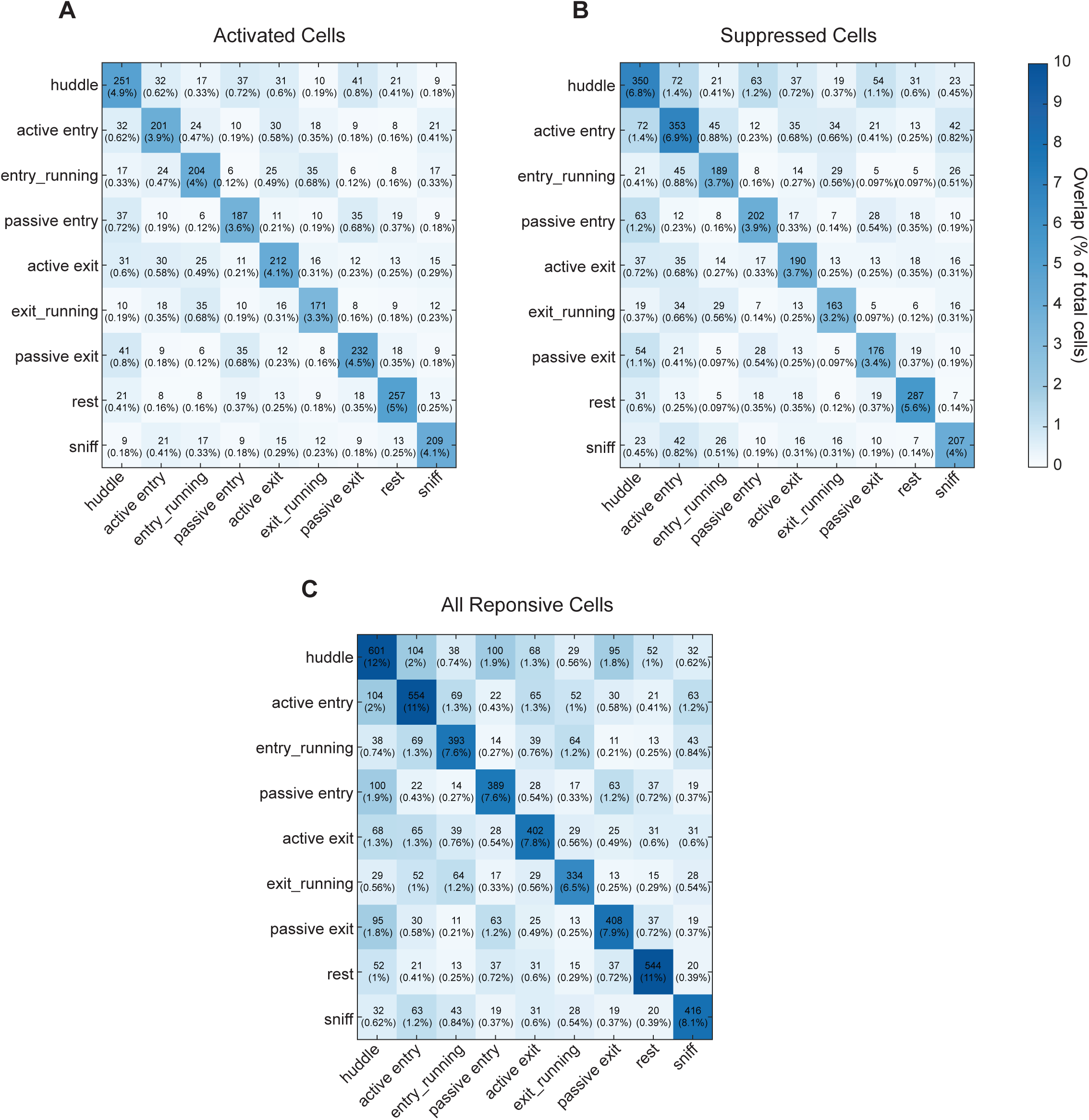
Matrices showing overlap of dmPFC cells responsive to various behaviors. **A.** Matrix showing number of cells activated by behaviors on x and y axis. Percentages correspond to percent of total imaged cells. **B.** Matrix showing number of cells suppressed by behaviors on x and y axis. Percentages correspond to percent of total imaged cells. **C.** Matrix showing number of all cells responsive to behaviors on x and y axis. Percentages correspond to percent of total imaged cells. Total # of imaged cells = 5141 from 11 animals.

**Supplementary Figure 8:**
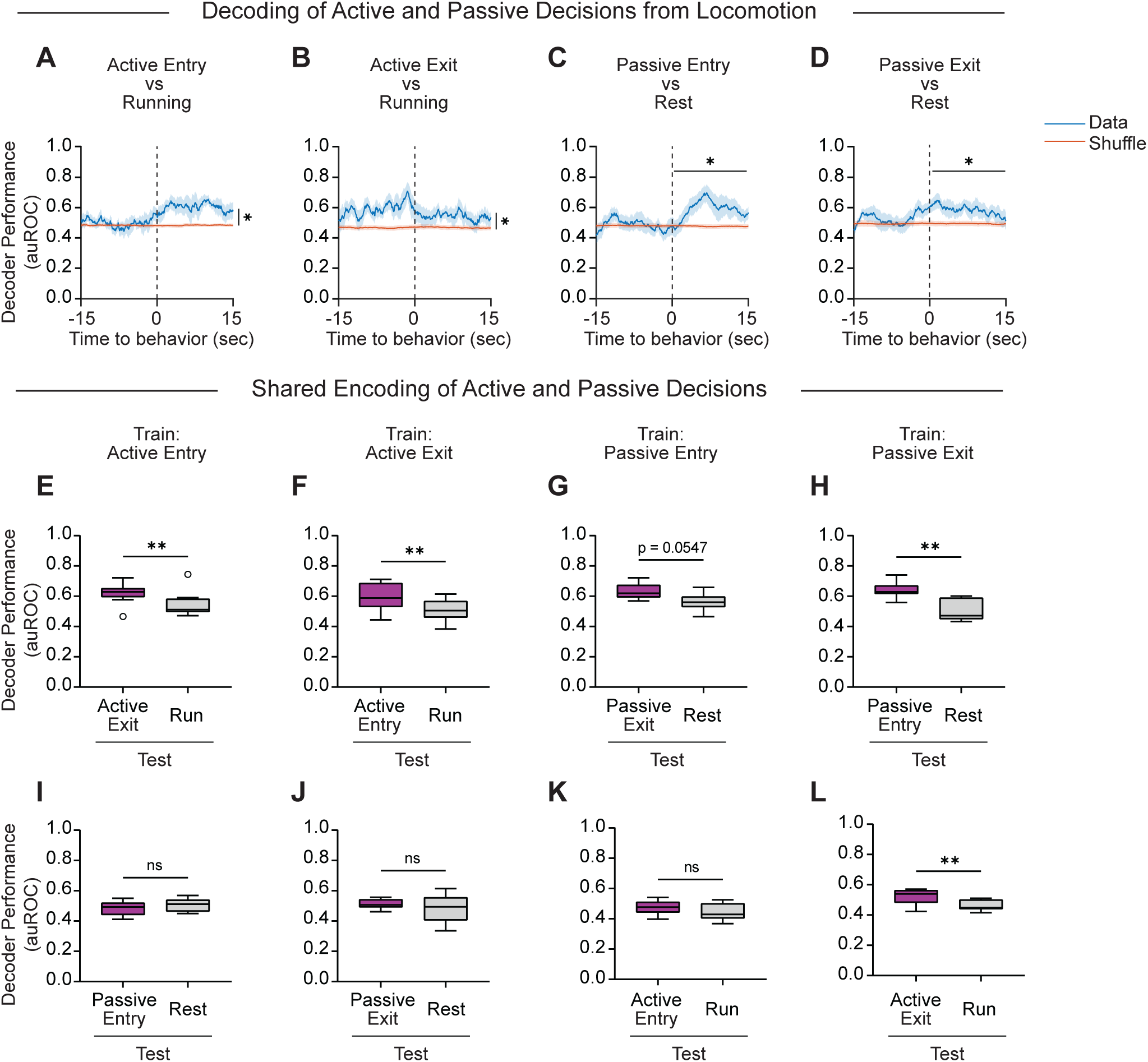
Additional dmPFC decoding of active and passive decisions. **A.** Performance of SVM decoders trained to classify active entry from speed-matched running. **B.** Performance of SVM decoders trained to classify active exit from speed-matched running. **C.** Performance of SVM decoders trained to classify passive entry from rest. **D.** Performance of SVM decoders trained to classify passive exit from rest. **E.** Performance of SVM decoders trained to classify active entry from baseline in predicting active exit from speed-matched running. **F.** Performance of SVM decoders trained to classify active exit from baseline in predicting active entry from speed-matched running. **G.** Performance of SVM decoders trained to classify passive entry from baseline in predicting passive exit from rest. **H.** Performance of SVM decoders trained to classify passive exit from baseline in predicting passive entry from rest. **I.** Performance of SVM decoders trained to classify active entry from baseline in predicting passive entry from rest. **J.** Performance of SVM decoders trained to classify active exit from baseline in predicting passive exit from rest. **K.** Performance of SVM decoders trained to classify passive entry from baseline in predicting active entry from speed-matched running. **L.** Performance of SVM decoders trained to classify passive exit from baseline in predicting active exit from speed-matched running. Box and whisker plots indicate the following: center line – median; box limits – upper and lower quartiles; whiskers – minimum and maximum values. Statistical tests include two-way repeated measures analysis of variance (ANOVA) with Bonferroni post-hoc tests (**A-D**) and Wilcoxon matched pairs tests (**E-L**). \**P*<.05, \*\**P*<.01, \*\*\**P*<.001, \*\*\*\**P*<.0001. See Supplementary Table 1 for details of statistical analyses.

**Supplementary Figure 9:**
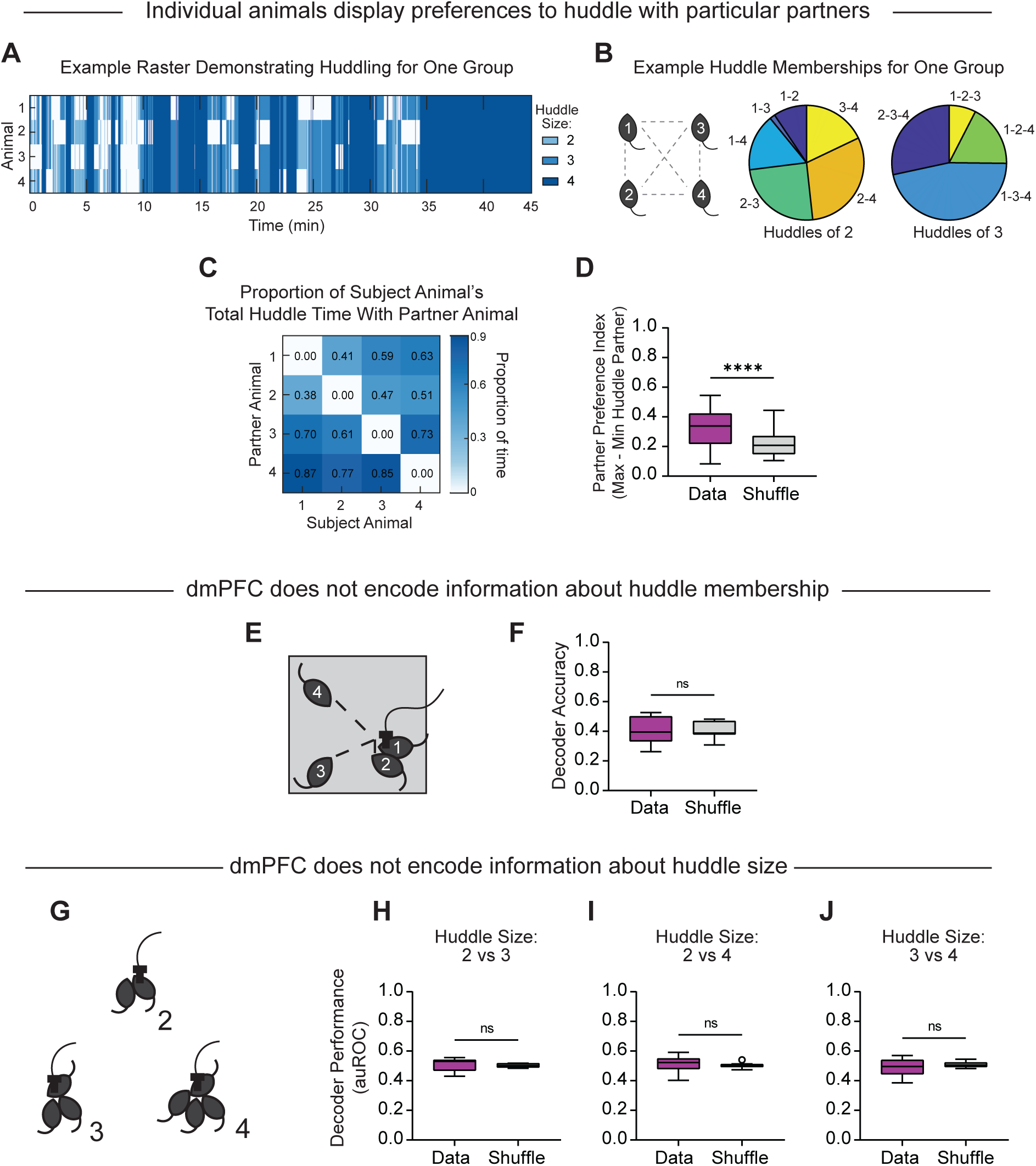
dmPFC does not encode huddle size or membership. **A.** Example raster plot demonstrating huddling behavior for all four animals in one session, color coded by huddle size. **B.** Example pie charts showing proportion of time for various huddle configurations for huddles of two and three for one group. **C.** Matrix demonstrating proportion of subject animal’s (x-axis) total huddle time with partner animals (y-axis) for one session. Sum of proportions for one animal can exceed 1 because subjects can huddle with more than one animal at a time in a larger huddle of two or three. **D.** Partner preference index (maximum preferred partner – minimum preferred partner) for real data versus a shuffled variation of the data in binary vectors containing individual huddle behaviors are circularly shifted relative to each other. **E.** Schematic illustrating potential huddle memberships for huddles of two during a miniscope imaging session. **F.** Performance of multi-class linear discriminant analysis (LDA) decoders trained to classify huddle membership for huddles of two from dmPFC population activity. Note that baseline is .33 because there are three possible memberships. **G.** Schematic illustrating potential huddle sizes during a miniscope imaging session. **H.** Performance of SVM decoders trained to classify huddle size of 2 from 3. **I.** Performance of SVM decoders trained to classify huddle size of 2 from 4. **J.** Performance of SVM decoders trained to classify huddle size of 3 from 4. Box and whisker plots indicate the following: center line – median; box limits – upper and lower quartiles; whiskers – minimum and maximum values. Statistical tests include Wilcoxon matched pairs tests (**D**,**F**,**H-J**). \**P*<.05, \*\**P*<.01, \*\*\**P*<.001, \*\*\*\**P*<.0001. See Supplementary Table 1 for details of statistical analyses.

**Supplementary Figure 10:**
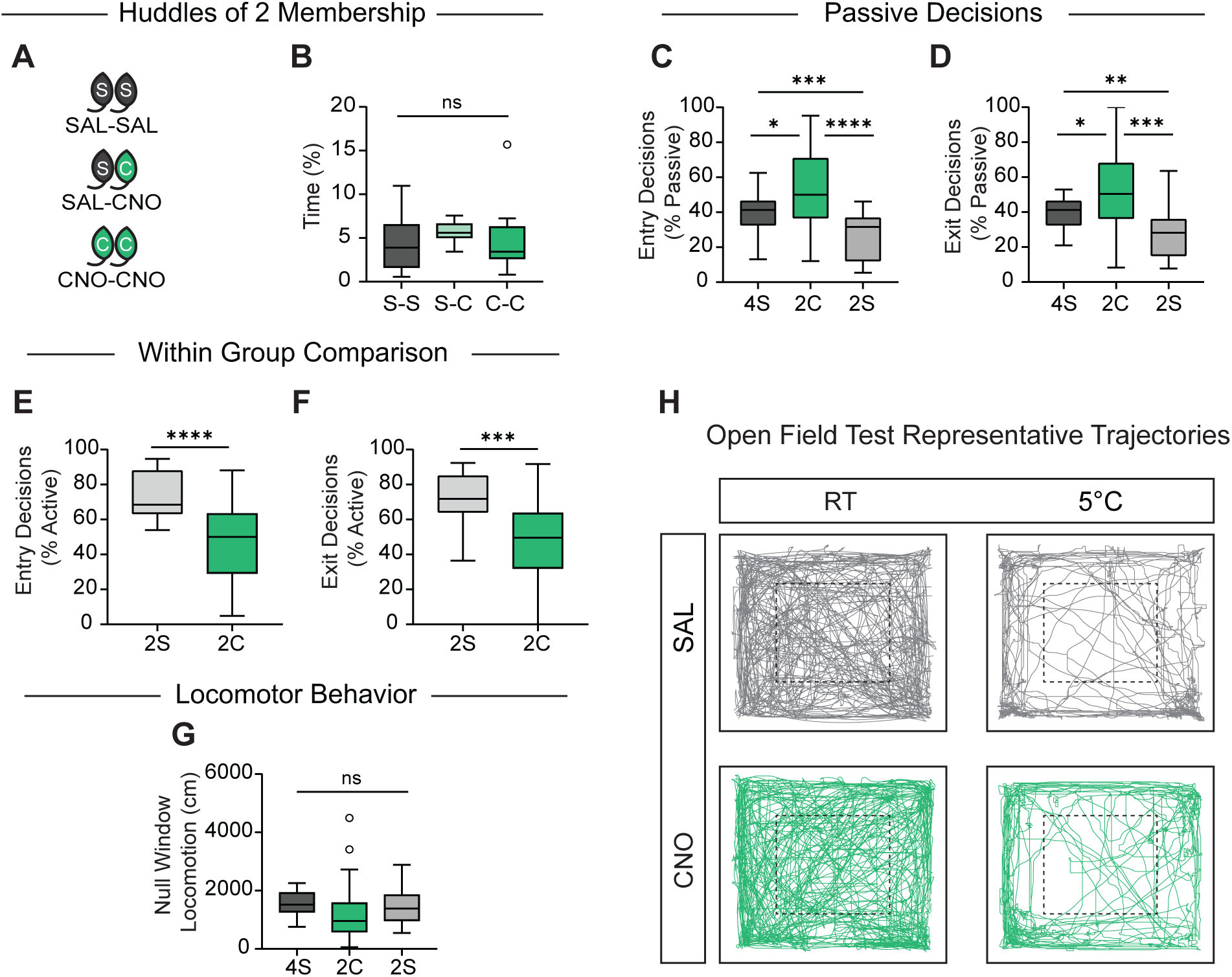
Additional data related to chemogenetic silencing in Figure 5. **A.** Schematic illustrating potential composition of membership for huddles of 2: SAL-SAL, SAL-CNO, and CNO-CNO during thermal challenge in 2C,2S condition. **B.** Percent of total time observed for possible membership compositions for huddles of two (n = 24 individuals from 6 groups). **C.** Within animal comparison of percent of entry decisions that are passive during 4S, 2C, and 2S conditions (n = 24 individuals from 6 groups). **D.** Within animal comparison of percent of exiting decisions that are passive during 4S, 2C, and 2S conditions (n = 24 individuals from 6 groups). **E.** Within group comparison of percent of entry decisions that are active for 2C, 2S condition (n = 24 individuals from 6 groups). Data in main figure shown as within animal comparisons. **F.** Within group comparison of percent of exiting decisions that are active for 2C, 2S condition (n = 24 individuals from 6 groups). Data in main figure shown as within animal comparisons. **G.** Individual animals’ total locomotion during null windows when no active, passive, or huddle behaviors are annotated during 4S, 2C, and 2S conditions (n = 24 individuals from 6 groups). **H.** Representative open field test trajectories at room temperature and 5°C after SAL or CNO injection. Box and whisker plots indicate the following: center line – median; box limits – upper and lower quartiles; whiskers – minimum and maximum values. Statistical tests include one-way repeated measures analysis of variance (ANOVA) with Bonferroni post-hoc tests (**B-D**,**G**) and Mann-Whitney tests (**E-F**). \**P*<.05, \*\**P*<.01, \*\*\**P*<.001, \*\*\*\**P*<.0001. See Supplementary Table 1 for details of statistical analyses.

**Supplementary Figure 11:**
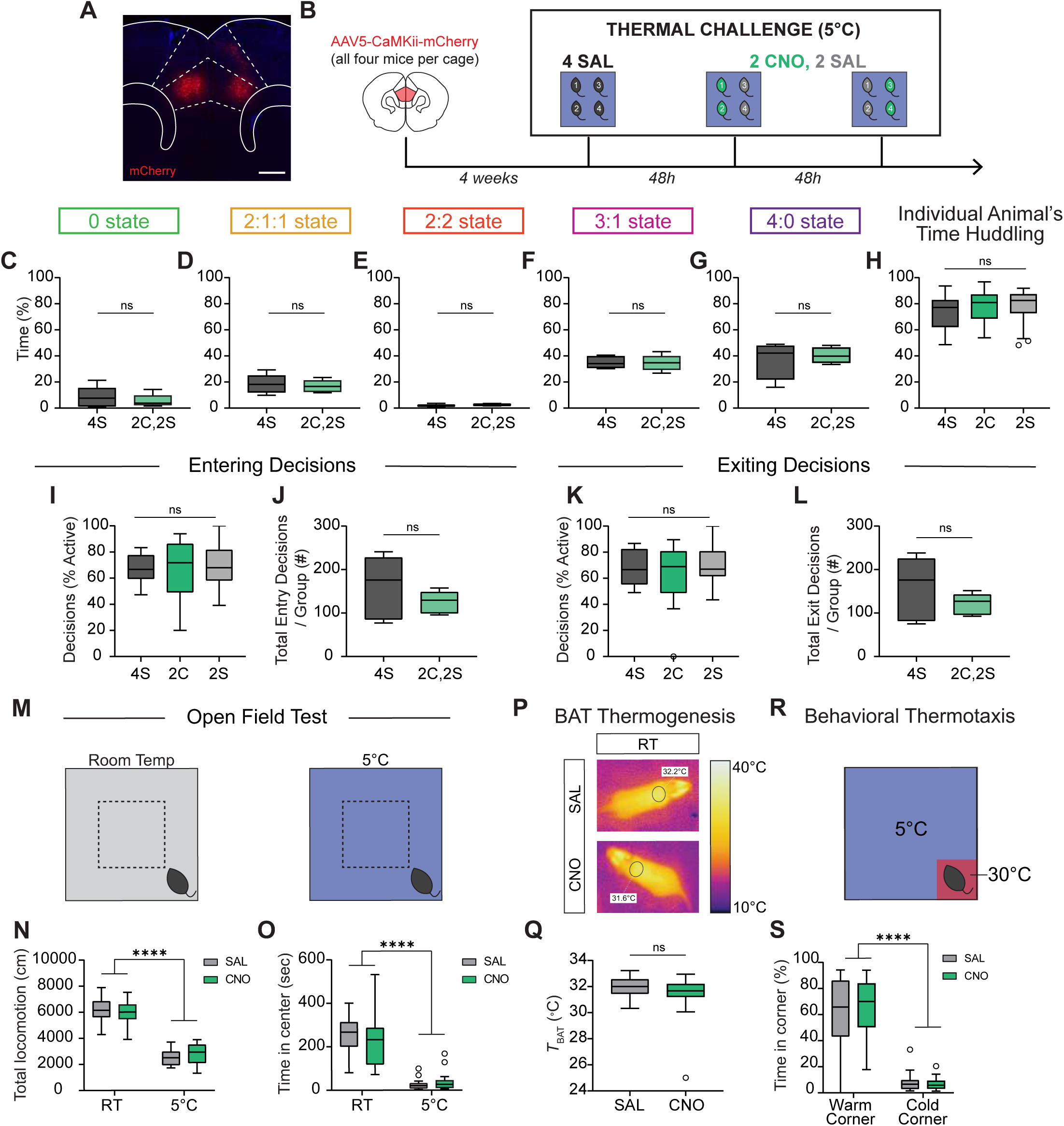
mCherry controls for chemogenetic silencing experiments. **A.** Example image showing AAV-mCherry expression in the dmPFC. Scale bar, 500 µm. **B.** Schematic illustrating experimental paradigm for mCherry chemogenetic control during thermal challenge. 4 SAL refers to condition in which all four animals are injected with saline. 2 CNO, 2 SAL refers to condition in which two animals are injected with CNO, and two with saline. **C-G.** Percent time in huddle states observed for all five group states during 4S and 2C,2S conditions (n = 5 groups). **H.** Individual animal’s total percent time spent huddling in 4S, 2C, and 2S conditions (n = 20 individuals from 5 groups). **I,K.** Within animal comparison of percent of entry or exit decisions that are active during 4S, 2C, and 2S conditions (n = 20 individuals from 5 groups). **J,L.** Total number of entry or exit decisions per group (active and passive from all four animals) during 4S and 2C,2S conditions (n = 20 individuals from 5 groups). **M.** Schematic illustrating open field test at room temperature (RT) and 5°C. **N.** Within animal comparison of total locomotion or time in center during open field test at both room temperature and 5°C after SAL or CNO injection (n = 20 animals). **P.** Representative infrared thermal images demonstrating temperature above BAT (brown adipose tissue, black circles) after SAL or CNO injection at room temperature. **Q.** Quantification of thermography images in regions above BAT after SAL or CNO injection (n = 20 animals). **R.** Schematic illustrating behavioral thermotaxis assay. **S.** Within animal comparison of percent time spent in warm corner versus the average of three cold corners after SAL or CNO injection (n = 20 animals). Box and whisker plots indicate the following: center line – median; box limits – upper and lower quartiles; whiskers – minimum and maximum values. Statistical tests include one-way (**H**,**I**,**K**) and two-way (**N**,**O**,**S**) repeated measures analysis of variance (ANOVA) with Bonferroni post-hoc tests, and Wilcoxon matched pairs tests (**C-G**,**J**,**L**,**Q**) tests. \**P*<.05, \*\**P*<.01, \*\*\**P*<.001, \*\*\*\**P*<.0001. See Supplementary Table 1 for details of statistical analyses.

## ACKNOWLEDGMENTS

We thank N. Ramesh for technical assistance, T. Pereira and the Pereira lab for assistance with SLEAP tracking, N. Sandoval for advice on thermal imaging, and all members of the Hong Laboratory for feedback and support. T.R. thanks F. Arbab for early conversations that inspired this research direction.

## FUNDING

NIH grant T32-NS048044 (TR)

NIH grant F32-MH123049 (TR)

NIH grant K99-MH133159 (TR)

Brain and Behavior Research Foundation Young Investigator Grant (TR)

NIH grant F31-MH134495 (KYL)

NIH grant R01-AG066821 (SMC)

NIH grant R01-MH130941 (WH)

R01-NS113124 (WH)

R01-MH132746 (WH)

RF1-NS132912 (WH)

Packard Fellowship in Science and Engineering (WH)

Vallee Scholar Award (WH)

Mallinckrodt Scholar Award (WH)

## COMPETING INTERESTS

The authors declare no competing interests.

## DATA AND MATERIALS AVAILABILITY

All data and code will be made available upon publication.

## SUPPLEMENTARY MATERIALS

Materials and Methods

Figs. S1 to S11

Supplementary Table 1

**Supplementary Table 1: Details of Statistical Analyses**

(see attached Excel File: Supplementary Table 1)

## MATERIALS AND METHODS

### Animals

All experiments were carried out in accordance with the National Institutes of Health Guide for Care and Use of Laboratory Animals and approved by the Institutional Animal Care and Use Committee (IACUC) of the University of California, Los Angeles. Subject mice were male and female C57BL6/J mice (stock number 000664) ordered from Jackson Laboratories at 8-12 weeks of age and 23-30g of weight. All mice were co-housed in groups of four animals. For calcium imaging, animals were cohoused in pairs separated by a perforated plastic acrylic divider for 3 weeks between lens implantation and baseplating. Following baseplating, animals were re-cohoused in groups of four with their original cage mates. Mice were maintained in a 12h:12h light/dark cycle (lighted hours: 10:00pm – 10:00 am) with *ad libitum* access to food and water. All experiments were performed during the dark cycle of the animals in a dark room with red or infrared light.

### Surgical Procedures for Miniscope and Chemogenetic Experiments

Mice were anaesthetized with 1.0 to 2.0% isoflurane and mounted on a stereotaxic device. All mice were given one subcutaneous injection of Ketoprofen (4mg/kg) on the day of surgery and Ibuprofen in drinking water (30mg/kg) starting on surgery day for 4-7 days. For Miniscope imaging experiments, viral injections and lens implantation in the dorsomedial prefrontal cortex (dmPFC) were performed. Specifically, we unilaterally injected 400-500 nL of AAV1-hSyn-GCaMP6f (Addgene #100837-AAV1) at 30 nL/min into the dmPFC using the stereotactic coordinates (AP: +2.0 mm, ML: ± .35 mm, DV: -1.8 mm). One week after injection, a 1.1mm diameter circular craniotomy was created (AP: +2.0, ML: .45 mm) and 1mm of tissue above the injection site was carefully aspirated using a 28-gauge needle. A 1.0 mm diameter GRIN lens (4.0 mm in length; Inscopix) was then implanted (AP: +2.0, ML: .45 mm, DV: -1.62 mm) and secured to the skull using dental cement and superglue. Three weeks after lens implantation, a baseplate and microscope were placed on top of the lens. The position and distance of the baseplate were adjusted using a Miniscope until the cells and blood vessels appeared sharp in the focal plane. The baseplate was then fixed to the skull at this distance using dental cement. Animals were handled and habituated to the Miniscope for 4-5 days prior to beginning behavior experiments.

For chemogenetic inhibition, AAV5-CaMKii-hM4Di-mCherry (Addgene #50477-AAV5) or AAV5-CaMKii-mCherry (Addgene #114469-AAV5) was injected. Given that dmPFC covers a relatively large area along the AP axis, we injected virus at two sites per hemisphere bilaterally to ensure effective inhibition. Specifically, we injected 300-350 nL of virus per site at AP +2.0 mm, ML: ± .4 mm, DV: -1.8 mm and AP +1.0 mm, ML: ± .4 mm, DV: -1.5 mm. Following surgeries, we allowed the mice to recover for 4 weeks before conducting behavioral testing.

### Histological Verification

After calcium imaging and chemogenetic experiments were completed, mice were transcardially perfused with 1X Phosphate Buffered Saline (PBS) and 4% paraformaldehyde (PFA). Brains were post-fixed in 4% PFA overnight at 4°C and cryo-protected for 48-72 hours in 30% sucrose at 4°C before freezing in OCT on dry ice. 35 um coronal sections were obtained using a cryostat (Leica Cm1950), and sections were stained with DAPI Fluoromount and mounted on Superfrost Plus Microscope Slides (Fisher). Images were acquired using an automated slide scanning fluorescence microscope (Leica DM6 B) to confirm the position of lens implantation and GCaMP6f virus expression for Miniscope experiments or hM4Di-mCherry virus for chemogenetic experiments.

### Huddling Behavior Assays

#### Thermal Challenge Assay

To assess group huddling behaviors, groups of four co-housed male mice were acclimated to behavioral box for 20 minutes per day for three days prior to onset of behavior testing. In addition, mice were habituated to human handling for 5 minutes. On the day of testing, animals were tail-marked with Sharpie pen in order to maintain identities when performing analyses post-hoc. Co-housed groups of four animals were placed in a 40 x 40 cm acrylic behavior box in a temperature controlled chamber (Thermo Scientific, PR505755R Refigerated Incubator) at 5°C to measure huddling behavior, or 20°C as a control, for 30-45 minutes, depending on the experiment. After behavioral testing, animals were returned to their home cage. For experiments testing titration of ambient temperature (Fig. S2, S3), animals were tested every 24 hours at 5°C, 10°C, 15°C, 20°C in scrambled order.

#### Huddling in Pairs versus Groups

To test differences in huddling behavior between pairs and groups of four animals, groups of four co-housed animals were tested under both conditions. Groups of four co-housed mice were acclimated to behavioral box for 20 minutes per day for three days prior to onset of behavior testing. In addition, mice were habituated to human handling for 5 minutes. On the day of testing, animals were tail-marked with Sharpie pen in order to maintain identities when performing analyses post-hoc. On the “pair” day, each group of four was split into two pairs, each of which was tested for 30 minutes. On the “group” day, all four animals were tested together. Pair and group conditions were tested 24 hours apart, counterbalanced by group size. Animals were placed in a 40 x 40 cm acrylic behavior box in a temperature controlled chamber at 5°C to measure huddling behavior for 30 minutes. After behavior testing, animals were returned to their home cage.

### Analysis of Animal Behavior

#### Development of Software for Multi-animal Behavior Annotation and Analysis

To label the behaviors of all 4 animals in our behavior experiments, we developed a Python application, BehaviorAnnotator. BehaviorAnnotator provides an interactive graphic interface and supports multiple streams of manual labeling for user-defined behaviors. Mainstream video formats are supported as well as the SEQ format files produced by specialized camera systems such as FLIR thermal camera or Point Grey camera. BehaviorAnnotator also displays multiple video streams in different layouts, allowing users to visualize camera recording from different angles to determine behavior labeling. BehaviorAnnotator is now available at Pypi (https://pypi.org/project/bannotator/) and can be easily installed through pip.

#### Pipeline for Analysis of Huddling Behavior

For thermal challenge huddling experiments, grayscale behavior videos were recorded at 30 frames per second (fps). Videos were subsequently analyzed through custom MATLAB scripts designed to identify huddling behaviors. Location, pose, and identity of the animals were automatically tracked using SLEAP (Social Leap Estimates Animal Poses, www.sleap.ai), an open-source neural network that we trained using manually labeled frames containing four animals interacting and/or huddling. For each frame, we extracted planar x and y coordinates for the animal’s nose, left ear, right ear, body center, and tailbase. Detection of huddling behavior incorporated two distinct analytical protocols: contour-based edge detection to identify huddle sizes, and SLEAP based identification of huddle membership. For contour-based edge detection, the video was filtered using a 3×3 blur kernel to binarize the pixels, complemented by using an “opening” image processing (erosion followed by dilation) using a 4×4 disk to remove fecal artifacts and tails. The background template was generated from the mean value of the first 10,000 frames, and binarization threshold was established at twice the standard deviation of the grayscale values. SLEAP predictions were then overlayed on top of extracted contours to map individual animals and determine instances of body contact. A temporal filter was applied to exclude contacts shorter than 2 seconds in duration. Automated detection of huddling behaviors was then visually checked by an experimenter, and manually annotated frame by frame to identify active and passive entry and exit events. Active and passive decisions are annotated as follows: active entry decisions are self-initiated behaviors that result in a change in behavior state from not huddling to huddling; passive entry decisions are behaviors initiated by other animals that result in a change in behavior state from not huddling to huddling; active exit decisions are self-initiated behaviors that result in a change in behavior state from huddling to not huddling; passive exit decisions are behaviors initiated by other animals that result in a change in behavior state from huddling to not huddling. If one animal active enters an existing huddling of two or three animals, the animals already in the huddle are not granted a passive entry, since they are already in the huddle. Likewise, if an animal actively exits a huddle, remaining animals are not granted a passive exit since they are still huddling.

For miniscope imaging experiments, no contouring process was applied due to interference from the miniscope cable. Instead, a method employing only SLEAP was developed, in which coordinates were mapped to box dimensions in millimeters and animal contacts were identified based on a threshold of 21mm, an optimal value derived from manual scoring. Videos were then scored for active and passive decisions, as well as for self-grooming, social investigation, and resting.

#### Superimposition of Huddles of 2

For superimposition of two animal huddling videos (Fig. 1), videos from huddling in pairs vs groups experiment (see Behavior Assays, above) were used. Videos and SLEAP coordinates were adjusted to the actual size of the enclosure using imwarp and transforPointsInverse functions. The adjusted videos were then superimposed onto each other frame by frame using four permutations: unrotated, 90 degree rotation, 180 degree rotation, and 270 degree rotation. These permutations control for the fact that in different videos, animals may opt to huddle in different corners of the enclosure. Edge detection (see Pipeline for Analysis of Huddling Behavior, above) and SLEAP predictions were used to assess the huddle states and membership. The rotation angle that led to the maximum huddling time was selected for comparison to the four animal huddling videos for behavior analysis.

### Chemogenetic Inhibition Experiments

#### Thermal Challenge Assay

Groups of co-housed mice were injected with AAV5-CaMKii-hM4di-mCherry, as described in Surgical Procedures above. Four weeks after surgeries, the mice were acclimated to the behavioral chambers following the protocol outlined in the Behavior Assays section above. In addition, mice were habituated to human handling and intraperitoneal injection by receiving injections of .25ml saline daily for 4 consecutive days. On the day of testing, mice were injected 30 minutes before behavior testing with either 1% dimethylsulfoxide (DMSO) in saline (as a control), or with clozapine-N-oxide (CNO) at 5 mg/kg body weight (Enzo, catalogue #BML-NS105) diluted in saline with 1% DMSO. Animals were returned to their home cage during the recovery period before behavior started. Animals were placed in a 40 x 40 cm acrylic behavior box in a temperature controlled chamber at 5°C for 30 minutes to measure huddling behavior. CNO and saline were administered per the following schedule: Day 1: all four animals received saline; Day 3: two animals received CNO, two animals received saline; Day 5: 2 CNO and 2 saline in a counterbalanced variation of Day 3; Day 7: all four animals received CNO.

#### Open Field Test

Groups of co-housed mice were injected with AAV5-CaMKii-hM4di-mCherry, as described in Surgical Procedures above. Four weeks after surgeries, the mice were acclimated to the behavioral box following the protocol outlined in the Behavior Assays section above. In addition, mice were habituated to human handling and intraperitoneal injection by receiving injections of .25ml saline daily for 4 consecutive days. On the day of testing, mice were injected 30 minutes before behavior testing with either 1% dimethylsulfoxide (DMSO) in saline (as a control), or with clozapine-N-oxide (CNO) at 5 mg/kg body weight (Enzo, catalogue #BML-NS105) diluted in saline with 1% DMSO. Animals were returned to a clean cage during the recovery period before behavior started. Animals were placed in a 40 x 40 cm acrylic behavior box in at 5°C or room temperature for 30 minutes. CNO and saline were administered 48 hours apart in a counterbalanced manner. Animal pose points were tracked using SLEAP (see Analysis of Animal Behavior, above). Time in center was defined as 25% of the total area, while periphery was defined as 75% of total area. The periphery consisted of the 10cm closest to the wall around the entire perimeter. Dependent measures were the overall ambulatory activity quantified as the total locomotion (in centimeters), and the time spent in the center (in seconds).

#### Measurement of Autonomic Thermoregulation

Groups of co-housed mice were injected with AAV5-CaMKii-hM4di-mCherry, as described in Surgical Procedures above. Four weeks after surgeries, the mice were acclimated to the behavioral box following the protocol outlined in the Behavior Assays section above. In addition, mice were habituated to human handling and intraperitoneal injection by receiving injections of .25ml saline daily for 4 consecutive days. On the day of testing, mice were first moved to a clean cage before injection and thermal acquisition began. Mice were injected with either 1% dimethylsulfoxide (DMSO) in saline (as a control), or with clozapine-N-oxide (CNO) at 5 mg/kg body weight (Enzo, catalogue #BML-NS105) diluted in saline with 1% DMSO. Infrared thermal images were captured using a thermal camera (FLIR One Pro LT) 60 minutes after CNO or saline injection. CNO and saline were administered 48 hours apart in a counterbalanced manner. Brown adipose tissue (BAT) temperature was analyzed using FLIR Ignite software. BAT skin temperature was the average temperature of a circular region above interscapular BAT.

#### Behavioral Thermoregulation Assay

Groups of co-housed mice were injected with AAV5-CaMKii-hM4di-mCherry, as described in Surgical Procedures above. Four weeks after surgeries, the mice were acclimated to the behavioral box following the protocol outlined in the Behavior Assays section above. In addition, mice were habituated to human handling and intraperitoneal injection by receiving injections of .25ml saline daily for 4 consecutive days. On the day of testing, mice were injected 30 minutes before behavior testing with either 1% dimethylsulfoxide (DMSO) in saline (as a control), or with clozapine-N-oxide (CNO) at 5 mg/kg body weight (Enzo, catalogue #BML-NS105) diluted in saline with 1% DMSO. Animals were returned to a clean cage during the recovery period before behavior started. Animals were placed in a 40 x 40 cm acrylic behavior box in at 5°C for 15 minutes. One corner of the box was warmed to 30°C by placing Hand Warmers (HotHands) on the outside of the box for 1 hour prior to behavior. CNO and saline were administered 48 hours apart in a counterbalanced manner. The corner used as a warm corner was also counterbalanced. Animal pose points were tracked using SLEAP (see Analysis of Animal Behavior, above), and the animal’s time spent in each corner was measured.

### Microendoscopic Calcium Imaging

Miniscope imaging experiments were carried out at least 7 days after base plates were affixed to the skull. Mice were habituated to human handling procedures and the microendoscope for 4-5 days prior to behavior, and habituated to the behavior box for at least 1 day before. Animals were outfitted with a customized microendoscope (UCLA Miniscope V4, purchased from OpenEphys; https://github.com/Aharoni-Lab/Miniscope-DAQ-QT-Software) and allowed to recover from handling for 10 minutes prior to behavior. One animal was recorded from per session, in the presence of three cagemate animals at 5°C. Calcium fluorescence videos and behavior videos were simultaneously recorded using UCLA Miniscope software at 15 frames per second. Animals were allowed to engage in huddling behavior for 30-45 minutes. Microendoscopes were connected to a digital acquisition device (DAQ) through a flexible, ultralight coaxial cable. Huddling behavior was automatically identified using the behavior analysis pipeline described above, and then manually checked and corrected frame by frame by a human annotator to identify onset and offset times of behaviors displayed by subject animals. Additional behaviors were also manually annotated (active entry, passive entry, active exit, passive exit, self-groom, social investigation).

### Extraction and Preprocessing of Calcium Signals

Raw videos from each imaging session were processed using an integrated Miniscope Analysis package. Briefly, raw videos were first processed for motion correction using the NormCorre algorithm (https://github.com/flatironinstitute/NoRMCorre). Motion corrected videos were then normalized using a custom written dF/F script in which *F-F_0_/F_0_* was applied to each frame, where F_0_ was the de-trended mean frame. dF/F normalized videos were de-noised using an FFT spatial band-pass filter in ImageJ (v1.53t, U.S. National Institutes of Health), and spatially down-sampled by a factor of 2 prior to ROI identification. We then identified putative cell bodies for extraction of neural signals using constrained non-negative matrix factorization (CNMF-E, https://github.com/zhoupc/CNMF_E) to isolate cellular signals and associated regions of interest. As CNMF-E can identify fluorescence changes from non-neuronal sources, such as motion artifacts or neuropil signals, traces from extracted cells were manually inspected to remove components that did not represent cell bodies, or to merge neighboring ROIs that came from one cell. Putative neurons that had abnormally shaped cell bodies (abnormally large or small), or that had calcium transients with low signal-to-noise were excluded from further analysis (< 5% of all putative neurons were excluded in this manner). dF/F calcium traces of individual cells were z-scored and presented in units of standard deviation (s.d.) before downstream analysis.

### Calcium Imaging Analysis

#### Single Cell Response Analyses

Single neuron’s responses to huddling and related behaviors are quantified using a receiver operating characteristic (ROC) analysis. For longer huddle bouts (more than 20 seconds), only the first 20 seconds of huddle are taken as the positive class, and the time perisods with no behavior were taken as the negative class. Prior to downstream analysis, all calcium traces were z-scored and presented throughout in units of standard deviation. We applied receiver operating characteristic (ROC) analysis to identify neurons that significantly responded during each type of investigation. We applied a binary threshold to the ΔF/F signal, classifying each time point as either indicating or not indicating a specific event. The true positive rate and false positive rate were computed over a range of binary thresholds that spanned the full range of the neural signal. These rates were used to construct an ROC curve, which depicts the detection capability of the neural signal at various thresholds. The area under the ROC curve (auROC) was then determined to quantify how strongly neural activity was influenced by each event. To evaluate significance, the observed auROC was compared against a null distribution, generated by circularly permuting the calcium signals with random circular time shifts 1000 times. A neuron was deemed significantly responsive (α < 0.05) if its auROC exceeded the 97.5th percentile (indicating activation) or fell below the 2.5th percentile (indicating suppression) of the null distribution. To control for the general encoding of locomotion when comparing active entry/exit behaviors, an equivalent number of speed-matched running bouts were identified using speed vectors calculated from the SLEAP tracked body locations. Running bouts are taken during periods with no annotated behaviors and when animals’ speed increases to the average speed of the corresponding entry/exit behaviors. For rest bouts, all animals were included in ROC analysis. For visualization of event-triggered averages in Figure 3D-E, only animals with a minimum of 5 rest bouts per session were included.

#### Principal Component Analysis

To visualize population responses during huddling behaviors, we applied principal component analysis (PCA) to obtain components that capture the covariance of the neural population during behavior events. Trial-averaged responses were computed over a time window of -5 to 10 seconds for each neuron per behavior event, and concatenated across event types (huddle, social investigation, rest; or active entry, passive entry, active exit, passive exit). Responses for each neuron were formed into a matrix which was used to perform PCA. Population vectors were then averaged over individual behavior bouts and projected onto the first 2 principal components for visualization. For comparison of population responses we calculated the Euclidean distances between PC-projected populations (using the first 2 principal components) within or across behaviors. To keep analyses across behaviors comparable, a maximum of 10 bouts per behavior per animal were used.

#### Population Decoding of Huddle, Social Investigation, and Rest

A support vector machine (SVM) decoder was trained to decode huddling from social investigation and speed-matched rest using the z-scored population calcium activities. For each imaging session, average neural activities of each behavioral bout are calculated for the two behavioral classes. For bouts that are longer than 10 seconds, the first 10 seconds are averaged. Bouts of the two behavioral classes are balanced by randomly drawing from the class with more bouts, such that the number of bouts are equal. Performance of the decoder performance is tested using a leave-one-out cross-validation (LOOCV) procedure, where one bout serves as the test set and the rest as the training set which is repeatedly tested for all bouts. To eliminate contamination, the training samples that are within 15 seconds from the test sample are eliminated from the training set. The test samples’ prediction scores are compared against the true labels to produce auROC. To generate the shuffled performance, calcium activities are circularly shifted with random time lag against the behaviors for 500 times, and an auROC is calculated for each shuffle. For each imaging session, the averaged auROC of 500 shuffles is compared to the averaged auROC of 50 runs from the experiment data. Animals that had less than 10 manually annotated rest bouts were supplemented with additional speed-matched immobility bouts identified using SLEAP data. For each analysis, animals that did not have a minimum of 10 behavior bouts were excluded.

#### Population Decoding of Active and Passive Decisions

A support vector machine (SVM) decoder was trained to decode active and passive decisions from each other using the z-scored population calcium activities. Active decisions and passive decisions are decoded using calcium activity along a 30-second time window centered at behavioral onset. Behaviors are first aligned from 15 seconds before onset to 15 seconds after onset and balanced with a random bootstrap, as described above. For each frame in the time series, an SVM decoder was trained on the z-scored population calcium activities of that frame in all behavioral bouts and tested using a leave-one-out cross-validation (LOOCV) procedure as described above. To generate the shuffled performance, calcium activities are circularly shifted with random time lag against the behaviors for 500 times, and an auROC is calculated for each shuffle. For each imaging session, the averaged auROC of 500 shuffles is compared to the averaged auROC of 50 runs from the experiment data. For each analysis, animals that did not have a minimum of 5 behavior bouts were excluded.

### Female Huddling Behavior

To assess group huddling behaviors, groups of four co-housed female mice were acclimated to behavioral box for 20 minutes per day for three days prior to onset of behavior testing. In addition, mice were habituated to human handling for 5 minutes. On the day of testing, animals were tail-marked with Sharpie pen in order to maintain identities when performing analyses post-hoc. Co-housed groups of four animals were placed in a 40 x 40 cm acrylic behavior box in a temperature controlled chamber (Thermo Scientific, PR505755R Refigerated Incubator) to measure huddling behavior for 45 minutes. After behavioral testing, animals were returned to their home cage. Animals were tested every 24 hours at 5°C, 10°C, 15°C, 20°C in scrambled order. Analysis of huddling behavior states was carried out in an identical fashion to male animals (see Pipeline for Analysis of Huddling Behavior in Main Methods). Fecal matter was quantified by counting the total number of feces on the last frame of each behavior video, for both males and females at 5°C and 20°C.

### Female and Male Thermotaxis Behavior

Age-matched adult male and female mice were acclimated to the behavioral box following the protocol outlined in the Behavior Assays section in the Main Methods. In addition, mice were habituated to human handling 5 minutes on each day. On the day of testing, animals were placed in a 40 x 40 cm acrylic behavior box in at 5°C for 15 minutes. One corner of the box was warmed to 30°C by placing Hand Warmers (HotHands) on the outside of the box for 1 hour prior to behavior. CNO and saline were administered 48 hours apart in a counterbalanced manner. The corner used as a warm corner was also counterbalanced. Animal pose points were tracked using SLEAP (see Analysis of Animal Behavior, above), and the animal’s time spent in each corner was measured.

### Support Vector Machine Decoding of Active and Passive Decisions from Locomotion

A support vector machine (SVM) decoder was trained to decode active decisions from speed-matched running bouts, and passive decisions from rest bouts, using the z-scored population calcium activities. Behavior classes are decoded using calcium activity along a 30-second time window centered at behavioral onset. Behaviors are first aligned from 15 seconds before onset to 15 seconds after onset and balanced with a random bootstrap, as described in the main methods. For each frame in the time series, an SVM decoder was trained on the z-scored population calcium activities of that frame in all behavioral bouts and tested using a leave-one-out cross-validation (LOOCV) procedure as described above. The decoder accuracy was compared with that of 500 random circularly shifted activities. Manually annotated rest bouts were supplemented with additional speed-matched immobility bouts identified using SLEAP data. For each analysis, animals that did not have a minimum of 5 behavior bouts were excluded.

### Mutual Decoding of Active and Passive Behaviors

Shared encoding of active and passive behaviors was assessed by training an SVM decoder on one behavior and testing on another. For instance, to test a decoder trained with active enter on active leave, a decoder is trained to decode active enter bouts against the same number of randomly drawn null bouts (no annotated behavior). The decoder is then tested to decode active leave versus randomly drawn null bouts. Performance is calculated as auROC of the test scores against true labels. Performance is compared to performance on a speed-matched running control when the test set is an active decision, and a rest control when the test set is a passive decision. Animals that had less than 10 manually annotated rest bouts were supplemented with additional speed-matched immobility bouts identified using SLEAP data. For each analysis, animals that did not have a minimum of 10 behavior bouts were excluded.

### Calculation of Partner Preference Index for Huddle Memberships

To assess animals’ preference to huddle with other group members, a preference index was calculated for each animal using the following equation: (T_max_ - T_min_)/T_total_. Where T_max_ is the total huddle time with the most preferred member, Tmin is the total huddle time with the least preferred member, and T_total_ is the total huddle time for the subject animal. The preference index is compared with a shuffle that controls for each animal’s total huddle time respectively. The binary vector representing each animal’s frame-by-frame huddle status are circularly shifted against each other to create temporal misalignment between animals. The time frames where only one animal is engaged in the huddle are randomly matched so that a huddle is composed of at least two animals. One thousand shuffles were created for each method and the averaged shuffle preference index was compared to true preference index.

### Multi-class Linear Discriminant Analysis (LDA) of Huddle Membership

A multi-class Linear Discriminant Analysis (LDA) was used to decode membership from calcium activities. For each experiment, huddle bouts when the imaged animal huddles with each of the 3 partners are averaged for decoding, each partner forming one class. For huddles longer than 10 seconds, only the first 10 seconds are averaged. Huddles for the 3 classes are balanced with random bootstrapping, trained with a 3-class linear discriminant analysis classifier and tested using a leave-one-out cross validation (LOOCV) procedure. The decoder accuracy was compared with that of 500 random circularly shifted activities. For each analysis, animals that did not have a minimum of 5 behavior bouts were excluded.

### Decoding of Huddle Size

A support vector machine (SVM) decoder was trained to decode huddles of 2, 3, and 4 in a pairwise manner using the z-scored population calcium activities. For each imaging session, average neural activities of each behavioral bout are calculated for the two behavioral classes. For bouts that are longer than 10 seconds, the first 10 seconds are averaged. Bouts of the two behavioral classes are balanced by randomly drawing from the class with more bouts, such that the number of bouts are equal. Performance of the decoder performance is tested using a leave-one-out cross-validation (LOOCV) procedure, where one bout serves as the test set and the rest as the training set which is repeatedly tested for all bouts. To eliminate contamination, the training samples that are within 15 seconds from the test sample are eliminated from the training set. The test samples’ prediction scores are compared against the true labels to produce auROC. To generate the shuffled performance, calcium activities are circularly shifted with random time lag against the behaviors for 500 times, and an auROC is calculated for each shuffle. For each imaging session, the averaged auROC of 500 shuffles is compared to the averaged auROC of 50 runs from the experiment data. For each analysis, animals that did not have a minimum of 10 behavior bouts were excluded.

### Social Preference Assay Behavior and Calcium Imaging

For calcium imaging during the social preference assay, animals were outfitted with the head-mounted Miniscope, briefly habituated in their home cage for 2-3 minutes, and then placed in a 40 x 40 cm arena. The arena contained two pencil wire cups in opposing corners. One cup contained an unfamiliar adult male, while the other contained an inanimate toy mouse. The subject animals were allowed to freely move about the environment and investigate social and toy stimuli at will for the duration of the 30 minute session. Subjects were imaged at room temperature and at 5°C 48 hours apart in a counterbalanced manner. We imaged 4937 neurons from 10 animals at room temperature and 4884 neurons from the same 10 animals at 5°C. Stimulus animals were habituated to the pencil cups for 20 minutes per day for 3 days prior to experiments. Subject animal pose points (nose, left ear, right ear, body, tail base) were tracked using SLEAP (see Main Methods). We considered investigation events to be periods where the animal’s head was within 3 inches of the center of the cup, and the angle between its head and the cup was < 60 degrees. Behavior annotations were converted into binary vectors that denote precisely which frames the animal is engaged in social vs toy investigation for downstream analysis.

### Social Preference Assay Single Cell Analysis

In the social preference assay, we analyzed the responses of individual dmPFC neurons when subjects closely investigated either the social or the toy chamber. Prior to downstream analysis, all calcium traces were z-scored and presented throughout in units of standard deviation. We applied receiver operating characteristic (ROC) analysis to identify neurons that significantly responded during each type of investigation. We applied a binary threshold to the ΔF/F signal, classifying each time point as either indicating or not indicating a specific event. The true positive rate and false positive rate were computed over a range of binary thresholds that spanned the full range of the neural signal. These rates were used to construct an ROC curve, which depicts the detection capability of the neural signal at various thresholds. The area under the ROC curve (auROC) was then determined to quantify how strongly neural activity was influenced by each event. To evaluate significance, the observed auROC was compared against a null distribution, generated by circularly permuting the calcium signals with random circular time shifts 2000 times. A neuron was deemed significantly responsive (α < 0.05) if its auROC exceeded the 97.5th percentile (indicating activation) or fell below the 2.5th percentile (indicating suppression) of the null distribution. This analysis included only time points marked as the behavior event of interest (social investigation or toy investigation) and baseline points which excludes all annotated events, ensuring that the identification of neurons responsive to a specific behavior was not confounded by their activity during other behaviors.

### Decoding of Social vs Toy Investigation During Social Preference Assay

To assess population level decoding of social vs object interaction from dmPFC neural data, we applied Linear Support vector machine (SVM) to identify hyperplanes that best separate the pair of population vectors associated with different events, using a leave-one-out prediction cross-validation (LOOCV) approach. We averaged the mean population activity associated with each independent event bout lasting at least 1.5 seconds. The mean activity for each cell was calculated over the entire bout duration, up to a maximum of 10 seconds post-event onset. The leave-one-out cross-validation method was then applied. For each test sample in each validation fold, we excluded samples within one minute before or after the test event onset to prevent temporal contamination between training and test datasets. we randomly down-sampled the majority class to match the minority class within the remaining training samples. The auROC value was computed for the predicted class probabilities. We generated shuffle controls by circularly shifting the events along the time axis 100 times to establish a chance level performance benchmark. These methods were applied to decode social investigation from toy investigation in a pairwise manner.

## Notes

### Competing Interest Statement

The authors have declared no competing interest.

## REFERENCES

1. Krause J, Ruxton GD. Living in groups. Published online 2002:210. Accessed July 15, 2024. https://global.oup.com/academic/product/living-in-groups-9780198508182

2. Sumpter D, Buhl J, Biro D, Couzin I. Information transfer in moving animal groups. Theory Biosci. 2008;127(2):177–186. doi:10.1007/S12064-008-0040-1

3. Couzin ID. Collective cognition in animal groups. Trends Cogn Sci. 2009;13(1):36–43. doi:10.1016/j.tics.2008.10.002

4. Berdahl A, Torney CJ, Ioannou CC, Faria JJ, Couzin ID. Emergent sensing of complex environments by mobile animal groups. Science (1979). 2013;339(6119):574–576. doi:10.1126/SCIENCE.1225883/SUPPL_FILE/574.MP3

5. Testard C, Shergold C, Acevedo-Ithier A, et al. Ecological disturbance alters the adaptive benefits of social ties. Science (1979). 2024;384(6702):1330–1335. doi:10.1126/SCIENCE.ADK0606

6. Sliwa J. Toward collective animal neuroscience. Science (1979). 2021;374(6566):397–398. doi:10.1126/SCIENCE.ABM3060/ASSET/6EAD86E1-E4E5-44AF-9461-DF7B76008B70/ASSETS/GRAPHIC/SCIENCE.ABM3060-F1.SVG

7. Yu JH, Napoli JL, Lovett-Barron M. Understanding collective behavior through neurobiology. Curr Opin Neurobiol. 2024;86:102866. doi:10.1016/J.CONB.2024.102866

8. Pereira TD, Tabris N, Matsliah A, et al. SLEAP: A deep learning system for multi-animal pose tracking. Nat Methods. 2022;19(4). doi:10.1038/s41592-022-01426-1

9. Chen Z, Zhang R, Fang HS, et al. AlphaTracker: a multi-animal tracking and behavioral analysis tool. Front Behav Neurosci. 2023;17:1111908. doi:10.3389/FNBEH.2023.1111908/BIBTEX

10. Lauer J, Zhou M, Ye S, et al. Multi-animal pose estimation, identification and tracking with DeepLabCut. Nature Methods 2022 19:4. 2022;19(4):496–504. doi:10.1038/s41592-022-01443-0

11. Kappel JM, Förster D, Slangewal K, et al. Visual recognition of social signals by a tectothalamic neural circuit. Nature 2022 608:7921. 2022;608(7921):146–152. doi:10.1038/s41586-022-04925-5

12. David Zada A, Schulze L, Yu JH, et al. Development of neural circuits for social motion perception in schooling fish. Current Biology. 2024;0(0). doi:10.1016/J.CUB.2024.06.049

13. Forli A, Yartsev MM. Hippocampal representation during collective spatial behaviour in bats. Nature 2023 621:7980. 2023;621(7980):796–803. doi:10.1038/s41586-023-06478-7

14. Rose MC, Styr B, Schmid TA, Elie JE, Yartsev MM. Cortical representation of group social communication in bats. Science (1979). 2021;374(6566). doi:10.1126/SCIENCE.ABA9584/SUPPL_FILE/SCIENCE.ABA9584_MDAR_REPRODUCIBILITY_CHECKLIST.PDF

15. Ramdya P, Lichocki P, Cruchet S, et al. Mechanosensory interactions drive collective behaviour in Drosophila. Nature 2014 519:7542. 2014;519(7542):233–236. doi:10.1038/nature14024

16. Ferreira CH, Moita MA. Behavioral and neuronal underpinnings of safety in numbers in fruit flies. Nature Communications 2020 11:1. 2020;11(1):1–10. doi:10.1038/s41467-020-17856-4

17. Harshaw C, Blumberg MS, Alberts JR. Thermoregulation, energetics, and behavior. APA handbook of comparative psychology: Basic concepts, methods, neural substrate, and behavior. Published online January 17, 2017:931–952. doi:10.1037/0000011-045

18. Research I of M (US) C on MN, Marriott BM, Carlson SJ. Physiology of Cold Exposure. Published online 1996. Accessed August 6, 2024. https://www.ncbi.nlm.nih.gov/books/NBK232852/

19. LeBlanc J. Mechanisms of adaptation to cold. Int J Sports Med. 1992;13 Suppl 1(SUPPL. 1). doi:10.1055/S-2007-1024629

20. El Marzouki H, Aboussaleh Y, Najimi M, Chigr F, Ahami A. Effect of Cold Stress on Neurobehavioral and Physiological Parameters in Rats. Front Physiol. 2021;12:660124. doi:10.3389/FPHYS.2021.660124/BIBTEX

21. Hu Y, Liu Y, Li S. Effect of Acute Cold Stress on Neuroethology in Mice and Establishment of Its Model. Animals (Basel). 2022;12(19). doi:10.3390/ANI12192671

22. Han A, Hudson-Paz C, Robinson BG, et al. Temperature-dependent differences in mouse gut motility are mediated by stress. Lab Animal 2024 53:6. 2024;53(6):148–159. doi:10.1038/s41684-024-01376-5

23. Kokolus KM, Capitano ML, Lee CT, et al. Baseline tumor growth and immune control in laboratory mice are significantly influenced by subthermoneutral housing temperature. Proc Natl Acad Sci U S A. 2013;110(50):20176–20181. doi:10.1073/PNAS.1304291110

24. Jung S, Lee M, Kim DY, Lee JW, Choe HK, Correspondence SYK. A forebrain neural substrate for behavioral thermoregulation. doi:10.1016/j.neuron.2021.09.039

25. Lal NK, Le P, Aggarwal S, et al. Xiphoid nucleus of the midline thalamus controls cold-induced food seeking. Nature 2023 621:7977. 2023;621(7977):138–145. doi:10.1038/s41586-023-06430-9

26. Yang S, Tan YL, Wu X, et al. An mPOA-ARCAgRP pathway modulates cold-evoked eating behavior. Cell Rep. 2021;36(6). doi:10.1016/J.CELREP.2021.109502

27. Glancy J, Groß R, Stone J V, Wilson SP. A Self-Organising Model of Thermoregulatory Huddling. PLoS Comput Biol. 2015;11(9):1004283. doi:10.1371/journal.pcbi.1004283

28. Alberts JR. Huddling by rat pups: Ontogeny of individual and group behavior. Dev Psychobiol. 2007;49(1):22–32. doi:10.1002/DEV.20190

29. Harshaw C, Culligan JJ, Alberts JR. Sex Differences in Thermogenesis Structure Behavior and Contact within Huddles of Infant Mice. PLoS One. 2014;9(1):e87405. doi:10.1371/JOURNAL.PONE.0087405

30. Harshaw C, Alberts JR. Group and individual regulation of physiology and behavior: A behavioral, thermographic, and acoustic study of mouse development. Physiol Behav. 2012;106(5):670–682. doi:10.1016/J.PHYSBEH.2012.05.002

31. Bautista A, García-Torres E, Martínez-Gómez M, Hudson R. Do newborn domestic rabbits Oryctolagus cuniculus compete for thermally advantageous positions in the litter huddle? Behav Ecol Sociobiol. 2008;62(3):331–339. doi:10.1007/S00265-007-0420-4

32. Harshaw C, Leffel JK, Alberts JR. Oxytocin and the warm outer glow: Thermoregulatory deficits cause huddling abnormalities in oxytocin-deficient mouse pups. Horm Behav. 2018;98. doi:10.1016/j.yhbeh.2017.12.007

33. Endo N, Ujita W, Fujiwara M, et al. Multiple animal positioning system shows that socially-reared mice influence the social proximity of isolation-reared cagemates. Communications Biology 2018 1:1. 2018;1(1):1–13. doi:10.1038/s42003-018-0213-5

34. Campbell LAD, Tkaczynski PJ, Lehmann J, Mouna M, Majolo B. Social thermoregulation as a potential mechanism linking sociality and fitness: Barbary macaques with more social partners form larger huddles. Scientific Reports 2018 8:1. 2018;8(1):1–8. doi:10.1038/s41598-018-24373-4

35. Ishizuka S. Do dominant monkeys gain more warmth? Number of physical contacts and spatial positions in huddles for male Japanese macaques in relation to dominance rank. Behavioural Processes. 2021;185:104317. doi:10.1016/J.BEPROC.2021.104317

36. Zitterbart DP, Wienecke B, Butler JP, Fabry B. Coordinated Movements Prevent Jamming in an Emperor Penguin Huddle. PLoS One. 2011;6(6):20260. doi:10.1371/journal.pone.0020260

37. Waters A, Blanchette F, Kim AD. Modeling Huddling Penguins. PLoS One. 2012;7(11). doi:10.1371/journal.pone.0050277

38. Canals M, Bozinovic F. Huddling behavior as critical phase transition triggered by low temperatures. Complexity. 2011;17(1):35–43. doi:10.1002/CPLX.20370

39. Schank JC, Alberts JR. Self-organized huddles of rat pups modeled by simple rules of individual behavior. J Theor Biol. 1997;189(1):11–25. doi:10.1006/jtbi.1997.0488

40. Báez-Mendoza R, Vázquez Y, Mastrobattista EP, Williams ZM. Neuronal Circuits for Social Decision-Making and Their Clinical Implications. Front Neurosci. 2021;15. doi:10.3389/fnins.2021.720294

41. Báez-Mendoza R, Mastrobattista EP, Wang AJ, Williams ZM. Social agent identity cells in the prefrontal cortex of interacting groups of primates. Science (1979). 2021;374(6566). doi:10.1126/SCIENCE.ABB4149

42. Padilla-Coreano N, Batra K, Patarino M, et al. Cortical ensembles orchestrate social competition through hypothalamic outputs. Nature. 2022;603(7902). doi:10.1038/s41586-022-04507-5

43. Kingsbury L, Huang S, Wang J, et al. Correlated Neural Activity and Encoding of Behavior across Brains of Socially Interacting Animals. Cell. 2019;178(2):429–446.e16. doi:10.1016/J.CELL.2019.05.022

44. Kingsbury L, Huang S, Raam T, et al. Cortical Representations of Conspecific Sex Shape Social Behavior. Neuron. 2020;107(5):941–953.e7. doi:10.1016/J.NEURON.2020.06.020

45. Chen P, Hong W. Neural Circuit Mechanisms of Social Behavior. Neuron. 2018;98(1):16–30. doi:10.1016/J.NEURON.2018.02.026

46. Gangopadhyay P, Chawla M, Dal Monte O, Chang SWC. Prefrontal-amygdala circuits in social decision-making. Nat Neurosci. 2021;24(1):5–18. doi:10.1038/S41593-020-00738-9

47. Li SW, Zeliger O, Strahs L, et al. Frontal neurons driving competitive behaviour and ecology of social groups. Nature. 2022;603(7902):661–666. doi:10.1038/S41586-021-04000-5

48. Yang J, Zhang H, Ni J, De Dreu CKW, Ma Y. Within-group synchronization in the prefrontal cortex associates with intergroup conflict. Nature Neuroscience 2020 23:6. 2020;23(6):754–760. doi:10.1038/s41593-020-0630-x

49. Zhang Z, Reis FMCV, He Y, et al. Estrogen-sensitive medial preoptic area neurons coordinate torpor in mice. Nature Communications 2020 11:1. 2020;11(1):1–14. doi:10.1038/s41467-020-20050-1

50. Cannon B, Nedergaard J. Nonshivering thermogenesis and its adequate measurement in metabolic studies. Journal of Experimental Biology. 2011;214(2):242–253. doi:10.1242/JEB.050989

51. Lim S, Honek J, Xue Y, et al. Cold-induced activation of brown adipose tissue and adipose angiogenesis in mice. Nat Protoc. 2012;7(3):606–615. doi:10.1038/NPROT.2012.013

52. Felix-Ortiz AC, Terrell JM, Gonzalez C, et al. Prefrontal Regulation of Safety Learning during Ethologically Relevant Thermal Threat. eNeuro. 2024;11(2). doi:10.1523/ENEURO.0140-23.2024

53. Grady F, Peltekian L, Iverson G, Geerling JC. Direct parabrachial-cortical connectivity. Cerebral Cortex. 2020;30(9). doi:10.1093/cercor/bhaa072

54. Tan CL, Cooke EK, Leib DE, et al. Warm-Sensitive Neurons that Control Body Temperature. Cell. 2016;167(1):47–59.e15. doi:10.1016/j.cell.2016.08.028

55. Landen J, Vandendoren M, Killmer S, Bedford N, Nelson A. Huddling substates in mice facilitate dynamic changes in body temperature and are modulated by Shank3b and Trpm8 mutation. Published online June 26, 2024. doi:10.21203/RS.3.RS-3904829/V1

56. Sotelo MI, Markunas C, Kudlak T, et al. Neurophysiological and behavioral synchronization in group-living and sleeping mice. Curr Biol. 2024;34(1):132–146.e5. doi:10.1016/J.CUB.2023.11.065

57. Kim J, Kim C, Han HB, et al. A bird’s-eye view of brain activity in socially interacting mice through mobile edge computing (MEC). Sci Adv. 2020;6(49). doi:10.1126/SCIADV.ABB9841/SUPPL_FILE/ABB9841_SM.PDF

58. Zhao Y, Yin X, Yu Y, et al. Social rank-dependent effects of testosterone on huddling strategies in mice. iScience. 2023;26(5). doi:10.1016/J.ISCI.2023.106516

